# Natural variation for unusual host responses and flagellin-mediated immunity against *Pseudomonas syringae* in genetically diverse tomato accessions

**DOI:** 10.1101/516617

**Authors:** Robyn Roberts, Samantha Mainiero, Adrian F. Powell, Alexander E. Liu, Kai Shi, Sarah R. Hind, Susan R. Strickler, Alan Collmer, Gregory B. Martin

## Abstract

The interaction between tomato and *Pseudomonas syringae* pv. tomato (*Pst*) is a well-developed model for investigating the molecular basis of the plant immune system. There is extensive natural variation in *Solanum lycopersicum* (tomato) but it has not been fully leveraged to enhance our understanding of the tomato-Pst pathosystem. We screened 216 genetically diverse accessions of cultivated tomato and a wild tomato species for natural variation in their response to three strains of *Pst*. The screen uncovered a broad range of previously unseen host symptoms in response to *Pst*, and one of these, stem galls, was found to be simply inherited. The screen also identified tomato accessions that showed enhanced responses to flagellin in bacterial population assays and in reactive oxygen species assays upon exposure to flagellin-derived peptides, flg22 and flgII-28. Reporter genes confirmed that the host responses were due primarily to pattern recognition receptor-immunity. This study revealed extensive natural variation in tomato for susceptibility and resistance to *Pst* and will enable elucidation of the molecular mechanisms underlying these host responses.

## Introduction

Bacterial speck is a persistent and economically important disease of tomato that causes necrotic lesions on leaves, stems, flowers, and fruits (Jones, 1991). The disease is caused by *Pseudomonas syringae* pv. tomato (*Pst*), which is a hemibiotrophic pathogen with a short epiphytic phase. *Pst* causes disease when it enters the apoplastic space of leaves through natural openings or wounds. When environmental conditions are favorable (cool, wet weather) and a susceptible host is available, *Pst* can multiply rapidly and eventually cause yellowing (chlorosis) and cell death (necrosis) when peak populations are reached.

The *Pst*/tomato pathosystem also serves as a model for investigating the molecular basis of bacterial pathogenesis and plant immunity (Pedley & Martin, 2003; Oh & Martin, 2011). *Pst* virulence is primarily determined by the suite of effectors that the pathogen translocates into plant cells via the type III secretion system (T3SS). The genome of DC3000, which is a widely-used strain of *Pst*, is 6.5 Mb and encodes 5,763 ORFs (Buell *et al*., 2003). In addition to 36 type III effectors (T3Es), which contribute to *Pst* virulence, DC3000 also produces a phytotoxin, coronatine (COR) that induces chlorosis and functions as a jasmonic acid-isoleucine mimic to promote colonization of plants by opening stomata and altering defense signaling pathways (Worley *et al*., 2013; Sakatos *et al*., 2018; Wei *et al*., 2018). By deleting 28 well-expressed T3Es and then reintroducing combinations of multiple effectors back into DC3000, eight effectors were found to be required for near wild-type *Pst* growth in *Nicotiana benthamiana* (Cunnac *et al*., 2011). The effectors AvrPto and AvrPtoB are two particularly well-studied T3Es which are major virulence determinants in DC3000 that greatly enhance bacterial growth in plant leaves (Lin & Martin, 2005; Cunnac *et al*., 2011).

Plants have evolved various mechanisms to defend themselves against pathogens. Constitutive defenses include many preformed barriers, such as cell walls, waxy cuticles, dense trichomes, and complex leaf topography/morphology. Defense responses can also be induced in plants through pattern recognition receptor (PRR)-triggered immunity (PTI) or nucleotide-binding oligomerization domain-like receptors (NLR)-triggered immunity (NTI). PTI is activated when plant PRRs recognize conserved microbe-associated molecular patterns (MAMPs). For example, the flagellin epitopes flg22 or flgII-28 are bound by the extracellular domains of leucine-rich repeat receptor kinases (LRR-RKs) FLAGELLIN SENSING 2 (FLS2; (Gomez-Gomez & Boller, 2000; Chinchilla *et al*., 2006; Sun *et al*., 2013)) or FLAGELLIN SENSING 3 (FLS3; (Hind *et al*., 2016)) respectively, and the csp22 epitope from bacterial cold shock proteins recognized by the LRR-RK cold shock protein receptor (CORE; (Wang *et al*., 2016)), while other unknown MAMPs such as one possibly recognized by Bti9 (Zeng *et al*., 2012) remain to be identified. Once a MAMP is recognized, a suite of host cellular events helps prevent microbial invasion and multiplication, including the rapid generation of reactive oxygen species (ROS), activation of MAPK cascades, callose deposition, calcium flux, stomatal closure, and transcriptional reprogramming (Monaghan & Zipfel, 2012; Li *et al*., 2016). PTI generally results in a moderate inhibition of pathogen growth (~10-fold), but has the potential to recognize more than one pathogen (Lacombe *et al*., 2010).

NTI activation is associated with localized programmed cell death (PCD), transcriptional reprogramming, MAPK cascade activation, ROS production, and significant inhibition of pathogen growth (~100 to 1000-fold). While NTI results in very effective suppression of pathogen growth and disease, it is usually pathogen- and strain-specific. In tomato, the only known NTI against *Pst* involves the recognition of the effectors AvrPto or AvrPtoB by the host intracellular Pto and Fen kinases acting with the NLR Prf (Pilowsky, 1982; Pedley & Martin, 2003; Rosebrock *et al*., 2007; Oh & Martin, 2011; Kraus *et al*., 2016). The presence of AvrPto and/or AvrPtoB define a *Pst* strain as race 0 whereas race 1 *Pst* strains lack these effectors and are not recognized by Pto/Prf. *Pto/Fen* and *Prf* occur within a 40-kb region on chromosome 5 in the wild relative *S. pimpinellifolium* and have been introgressed into many tomato varieties (Pilowsky, 1982; Pilowsky & Zutra, 1986; Martin *et al*., 1991; Martin *et al*., 1993). There are 12 species of wild relatives of tomato that are native to western South America, but only two wild species, *S. pimpinellifolium* and *S. habrochaites*, are known to have a functional Pto kinase that recognizes AvrPto and AvrPtoB (Martin *et al*., 1993; Riely & Martin, 2001; Rose *et al*., 2005; Peralta *et al*., 2008; Bao *et al*., 2015). Fen, which is in the same kinase family as Pto, encodes a protein that binds to the N-terminus of AvrPtoB and activates NTI in conjunction with Prf in the absence of the AvrPtoB C-terminal E3 ligase domain (Abramovitch *et al*., 2003; Abramovitch *et al*., 2006; Janjusevic *et al*., 2006; Rosebrock *et al*., 2007; Cheng *et al*., 2011; Mathieu *et al*., 2014). In the *S. chmielewskii* accession LA2677, the Pto protein recognizes AvrPtoB and not AvrPto and varies from the *S. pimpinellifolium* Pto by 14 amino acids (Kraus *et al*., 2016).

While Pto/Prf-mediated resistance to speck disease has been effective for decades, worldwide populations of *Pst* have shifted towards race 1 strains that evade Pto/Prf immunity through the loss of *avrPto/avrPtoB*, the failure to express AvrPtoB protein, or the expression of AvrPto/AvrPtoB variants that cannot be recognized by Pto (Lin *et al*., 2006; Kunkeaw *et al*., 2010; Cai *et al*., 2011). Race 1 strains were first detected in the United States in 1993 and have become increasingly common among strains found in field-grown tomatoes (Arredondo & Davis, 2000; Kunkeaw *et al*., 2010). There are three recent reports of resistance to race 1 *Pst* strains although in each case the resistance appears to be quantitative and its usefulness in controlling speck disease in the field will probably be limited (Kunkeaw *et al*., 2010; Cai *et al*., 2011; Bao *et al*., 2015). *Pst* strains with features intermediate between race 0 and race 1 strains have also been found that express AvrPto that is recognized by Pto, but appear to have another unknown factor that compromises Pto-mediated resistance (Kraus *et al*., 2017).

Probing of the host response to distinguish PTI and NTI responses is facilitated by the availability of *Pst* DC3000 variants that, for example, carry a deletion of the *fliC* gene which encodes flagellin (Kvitko *et al*., 2009; Rosli *et al*., 2013; Wei *et al*., 2013), or deletions of specific effector genes (Lin & Martin, 2005; Cunnac *et al*., 2011; Rosli *et al*., 2013; Liu *et al*., 2014; Wei *et al*., 2015). By comparing the host response to DC3000 versus DC3000Δ*fliC*, PTI resistance linked to flagellin recognition can be inferred. For NTI, DC3000 can be compared to DC3000Δ*avrPto*Δ*avrPtoB* to identify plants with Pto-mediated resistance or with *Pst* strains with a much different effector repertoire to reveal other novel NTI recognition mechanisms.

Here, we describe a screen of genetically diverse accessions of cultivated tomato and a wild tomato species to identify new host responses to *Pst*. We generated a phylogenetic tree based on genome sequence data from ~1,400 tomato accessions and identified a representative subset of 216 accessions comprised of cultivated tomato (*S. lycopersicum* and *S. lycopersicum* var. cerasiforme) and its most closely related wild species (*S. pimpinellifolium*). We spray inoculated the accessions with three *Pst* strains chosen to distinguish between PTI or NTI responses, and used bacterial growth assays, reporter genes, and ROS assays to identify accessions with unique PTI responses. We discovered accessions that developed previously unseen symptoms in response to *Pst*, and we examined the genetic basis of one of these, stem galls. These results reveal extensive novel natural variation in the response of tomato to *Pst* and lay the foundation for the identification and characterization of new plant genes involved in disease susceptibility and resistance.

## Materials and Methods

### Phylogenetic tree

Publicly available datasets (Sim *et al*., 2012; Lin *et al*., 2014; Tomato Genome Sequencing Consortium, 2014) were combined to obtain genome-wide base calls from 6,225 sites in tomato genome assembly version SL2.40 (Tomato Genome Consortium, 2012). Genotypes were converted to aligned fasta format and a maximum likelihood tree was generated with 100 bootstrap replicates using Mega version 7.0.16 (Sade-Feldman *et al*., 2017). The alignment and resulting tree were analyzed with ClusterPicker version 1.2.3 (Ragonnet-Cronin *et al*., 2013) with the following parameters: Initial support threshold=0.9, Support threshold=0.9, Genetic distance threshold=0.045, Large cluster threshold=10. The tree was visualized in FigTree version 1.4.2 (Rambaut, 2007). Reference allele frequency was plotted using Circos version 0.69-6 (Krzywinski *et al*., 2009).

### Plant growth conditions

*S. lycopersicum* (tomato) or *S. pimpinellifolium* seedlings were grown in Cornell Plus Mix soil in a greenhouse 24°C day/22°C night without supplemental lighting for 3 weeks before spray inoculation or 4 weeks before vacuum infiltration. For seed sources of each accession, see Table S2.

### Spray inoculations and disease screening

*Pseudomonas syringae* pv. tomato (*Pst*) strains were grown on King’s B medium with the appropriate antibiotic for 48 hours at 30°C. For each strain, the bacterial lawns were scraped from the plate, suspended in 10 mM MgCl_2_, and centrifuged for 10 minutes at 4000 rpm. The pelleted bacteria were resuspended in fresh 10 mM MgCl_2_ to 1 × 10^8^ CFU/mL (OD_600_ of 0.4 or 0.42 for the DC3000 strains or Pst25, respectively) with 0.04% Silwet L-77.

The initial spray inoculation screen was divided into seven experiments, with 30 accessions and the six controls in each experiment. Three plants of 30 accessions were included in every experiment. 16 hours prior to inoculation, the three-week-old seedlings were placed in a transparent box designed to hold 100% humidity. Separate plants were sprayed with each *Pst* strain on both the abaxial and adaxial sides of the leaves until runoff, and then placed back inside the box along with a HOBO^®^ data logger (Onset Computer Corporation) to monitor temperature, relative humidity, and light conditions inside the box (24°C day/22°C night, 100% relative humidity, 8 hour day length, and 150 μmol m^−2^ s^−1^ day/ 0 μmol m^−2^ s^−1^ night photosynthetic photon flux density (PPFD)). Host responses were scored at 3- and 6- dpi. Selected accessions were included together in a follow-up experiment in the same conditions listed above to confirm phenotypes.

### Disease scoring

Plants were blindly scored 3- and 6-days post inoculation (dpi) using a 1-5 disease rating scale. Scores from 6 dpi were averaged between the three plants for each accession. To normalize the scores between accessions in the initial screen, which was conducted over seven experiments, the average score of each of the six control accessions were calculated across the seven experiments. The averages of the control accessions in each experiment were calculated and divided by the overall averages of the six accessions combined to get the normalization factor. The average scores for the accessions in each experiment were multiplied by the normalization factor calculated for each experiment to normalize the scores. Normalization was not used in the follow-up experiment because all plants were screened together.

### Bacterial population assays

*Pst* strains were prepared for vacuum infiltration as described previously (Kraus *et al*., 2017) or spray inoculated as described above. Bacteria were suspended to a final concentration of 2 × 10^4^ CFU/mL (vacuum infiltrations) or 5 × 10^8^ CFU/mL (spray inoculations) in 10mM MgCl_2_. For the spray inoculated plants, leaflets were surface sterilized with half-strength bleach for 2 minutes prior to sampling. For both spray inoculated and vacuum infiltrated plants, bacterial populations were quantified as described by (Kraus *et al*., 2017). All experiments were repeated independently three times, and data shown are from one representative experiment (n=3 plants per accession per strain). Error bars represent the mean population of three plants +/− S.D. Significance was determined using a pairwise t-test and analyzed using the Prism 7 program (GraphPad Software).

To determine whether bacteria occur within stem galls, galls were excised from plant stems inoculated with DC3000Δ*avrPto*Δ*avrPtoB* using a razor blade and surface sterilized in 10% bleach for 2 minutes. The tissue was macerated and plated on King’s B medium supplemented with rifampicin, which is the antibiotic marker for the bacterial strain. Bacterial growth was observed two days after plating the tissue.

### Reactive oxygen species bioassays

ROS production in response to 100 nM of DC3000 flg22 or flgII-28 peptides was measured as previously described (Hind *et al*., 2016), and the average ROS response for each plant is the mean of three replicate leaf discs from two plants. The assay was repeated on each accession in at least two independent experiments. One representative experiment is shown in the figure.

### Reporter genes

Three, four-week-old plants were vacuum infiltrated with 5 × 10^6^ CFU/mL of *Pst*. Six hours post inoculation, three independent biological replicates were sampled from different plants inoculated with each strain. RNA extraction and qRT-PCR for the reporter genes were conducted as previously described (Pombo *et al*., 2014). Significance was determined using a pairwise t-test and was analyzed using the Prism 7 program (GraphPad Software).

### Gall imaging

Stem galls from LA1589 plants sprayed with 1 × 10^8^ CFU/mL DC3000Δ*avrPto*Δ*avrPtoB* were transverse-sectioned with a razor blade at 5 dpi and imaged with a Leica M205 stereoscope under darkfield conditions.

## Results

To choose accessions that represents the genetic diversity of cultivated tomato and a closely-related species, we developed a phylogenetic tree based on 6,225 SNPs derived from genome sequences of 1,429 accessions from the ‘SolCap,’ tomato ‘360,’ and tomato ‘150’ databases (Sim *et al*., 2012; Lin *et al*., 2014; Tomato Genome Sequencing Consortium, 2014) (Fig. S1 and Table S1). From the 83 clusters generated in this process, 181 accessions were chosen to represent the genetic diversity (Table S2). More accessions were included from larger clusters, and accession choices were also based on availability of seed. Each accession was given an index number based on its cluster (C01 - C83) and a letter identifier (A-Z) (Table S2). An additional 35 accessions were included that are of interest to various breeding programs (Table S2, CXA-ZZ) or are a parent in a tomato MAGIC population (Pascual et al 2014) (Table S2, MC001-8). In total, 216 accessions were included in the screen.

We spray inoculated each of the 216 accessions with: 1) *Pst* NY15125 (Pst25), a race 0 strain collected in New York that expresses AvrPto (Kraus et al 2017); 2) DC3000Δ*avrPto*Δ*avrPtoB* (DC3000ΔΔ); or 3) DC3000Δ*avrPto*Δ*avrPtoB*Δ*fliC* (DC3000ΔΔΔ) (Table 1). Different symptoms on an accession in response to the two DC3000 strains is potentially indicative of natural variation in flagellin-mediated PTI whereas different symptoms in response to DC3000ΔΔ and Pst25 is suggestive of variation in NTI in response to one of 25 TTEs that differ between these two strains (Table 1).

**Table 1.** *Pseudomonas syringae* pv. tomato strains used in this study. Salient features of each strain are shown, along with the strain used for comparison to hypothesize NTI, PTI, or other immune responses. other tested *S. pimpinellifolium* accessions.

| Strain | Features | Rationale | Compare to | Expected immune response | Reference |
| --- | --- | --- | --- | --- | --- |
| NY15125 (Pst25) | Isolated in NY. Has 27 effectors, including AvrPto (race 0). 12 are different from DC3000 | Identify novel NTI loci | DC3000ΔΔ | NTI | Kraus *et al.* (2017)<br>Mazo-Molina *et al.*, (2019) |
| DC3000<br>ΔavrPtoΔavrPtoB<br>(DC3000ΔΔ) | Has 28 effectors with 13 being different from Pst25. *avrPto* and *avrPtoB* deleted | Identify novel NTI loci, compare with DC3000ΔΔΔ for PTI | Pst25 | NTI | Lin *et al.* (2005) |
|  |  |  | DC3000ΔΔΔ | PTI |  |
| DC3000<br>ΔavrPtoΔavrPtoBΔfliC<br>(DC3000ΔΔΔ) | *avrPto*, *avrPtoB*, and *fliC* deleted | Compare with DC3000ΔΔ for PTI | DC3000ΔΔ | PTI | Kvitko *et al.* (2009) |

The three *Pst* strains were inoculated onto three plants of each accession, with 30 accessions being tested in each experiment. As controls, we used Rio Grande-PtoR (has the *Pto* gene), Rio Grande-prf3 (has a non-functional Pto pathway), Heinz 1706 (origin of the tomato reference genome), Moneymaker (commonly used in research), and two lines in the Rio Grande background with decreased expression of immunity-related genes: hpBti9 (PTI-related knockdown; (Zeng *et al*., 2012)) and hpPto (NTI-related knockdown; (Pascuzzi, 2006)). We inoculated the plants under conditions that maintained 100% humidity to aid pathogen infection. Each plant was scored on a 1-5 scale and scores were averaged between the three plants of the same accession (Fig. S2a,b). We screened all 216 accessions in seven experiments and normalized the scores of all the accessions among experiments based on the controls in order to compare the accessions (Table S3). The distribution of the scores among the accessions is shown in Fig. S2b. Accessions with interesting responses from this initial screen were repeated in a subsequent round of screening where all accessions and control lines could be scored in a single experiment and compared to the normalized scores of the first screen (Table S3).

### Some accessions developed unusual symptoms in response to *Pst* strains

In addition to scoring plants for overall disease, we recorded interesting and/or unusual host responses observed in the screen. We observed many accessions that displayed typical speck symptoms such as leaf specks with or without yellows halos (Fig. S2c and Tables S4-6). However, we also observed a remarkable array of other phenotypes that are not typically associated with *Pst* (Fig. 1 and Tables S4-6). While we typically observe specks on stems when plants are vacuum infiltrated, we more rarely saw specks on stems when plants were spray inoculated which is more similar to natural infection. Some accessions developed ‘shot holes’, in which the dry lesions dropped out of the leaf; we have not observed this before with bacterial speck, though it is known to occur in various bacterial and fungal diseases in many crop species (such as wildfire disease of tobacco, angular leaf spot of cucumber, and bacterial spot of stone fruit). We also observed necrotic spots, severe necrosis, chlorosis (large areas of yellowing), water soaking, and severe leaf abscission, which are not typically associated with bacterial speck disease. We observed raised, rough growths (‘galls’) on the stems of some accessions which we believe is a first observation for *Pst*. Interestingly, the galls were primarily found upon inoculation with the DC3000 strains and not with Pst25, suggesting there is a *Pst* strain-specific factor involved. Such strain-specificity was not observed for any of the other unusual phenotypes. Selected accessions that represent the diverse phenotypes observed, along with their disease scores and hypothesized NTI/PTI responses, are shown in Table 2. We also observed on some accessions that the high humidity alone caused unusual symptoms that were unrelated to the inoculation (Fig. S3). These symptoms included hypertrophic lesions on leaves and petioles and browning on the leaf margins and these accessions were not pursued further.

**Fig. 1.**
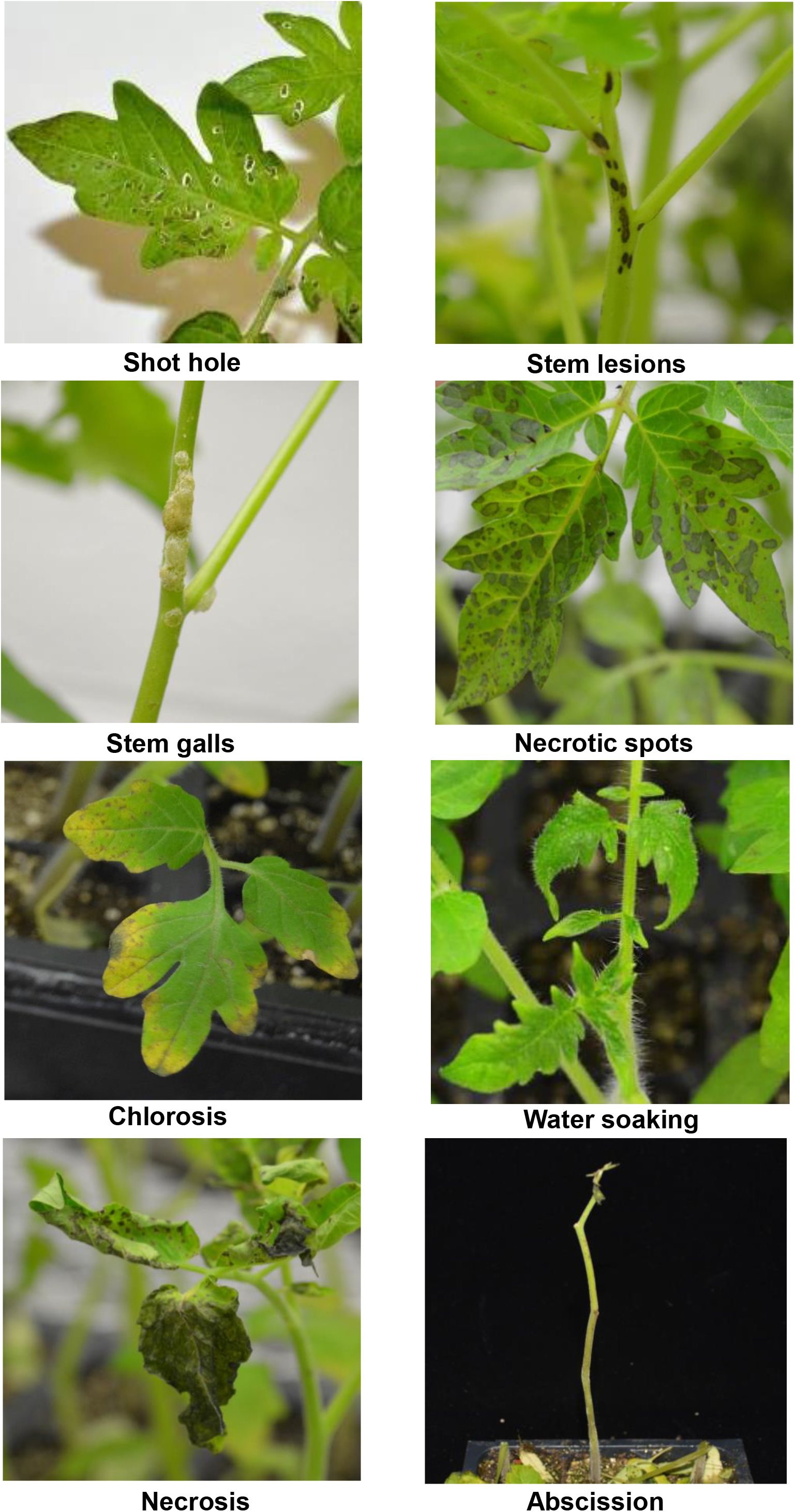
Unusual host responses after spray inoculating with *Pst*. Photos are representative of the unusual host responses observed in the screen.

**Table 2.** Tomato accessions that represent the various phenotypes observed in the screen. The averaged normalized scores are shown from the two independently repeated screens. Controls are shown at the bottom of the table and are the averaged scores from the original seven individual experiments, and were used to normalize the scores in the first replicate of the screen. In both screens, three plants of each accession were rated and the scores were averaged among the plants. Phenotypes are listed for each accession if they were observed and include: leaf specks (spk), chlorosis (chl), shot holes (sht), severe necrosis (nec), necrotic spots (spt), stem specks (stm), water soaking (wso), or stem galls (gal). Controls: Rio Grande-PtoR (RG-PtoR, NTI resistance), Rio Grande-prf3 (RG-prf3, susceptible), Heinz 1706 (reference genome, susceptible), Moneymaker (susceptible), Rio Grande-prf3-hairpinBti9 (hpBti9, has Bti9 knocked down using a hairpin construct, susceptible), and Rio Grande-PtoR-hairpinPto (hpPto, has Pto knocked down using a hairpin construct, susceptible).

| Index | Accession | Pst25 |  | DC3000ΔΔ |  | DC3000ΔΔΔ |  |
| --- | --- | --- | --- | --- | --- | --- | --- |
|  |  | Average score | Phenotype | Average score | Phenotype | Average score | Phenotype |
| C01B | EA02304 | 3.2 | spk | 2.7 | spk, spt | 3.9 | sht, spk, nec, chl |
| C03A | LA1511 | 2.8 | chl, spk | 1.9 | spk | 2.2 | gal, sht, stm, spk |
| C12A | Marmande VFA | 3.9 | spk | 2.9 | spk, chl | 4.0 | stm, spk, spt, nec |
| C13E | EA01356 | 3.1 | spk | 2.5 | sht, stm, spk | 3.6 | spk, chl |
| C21O | NC EBR-1 | 2.9 | spk, chl | 1.9 | spk | 2.5 | spk |
| C27A | M82 | 3.8 | spk | 2.2 | spk | 4.0 | gal, stm, spk |
| C41C | CR149 | 3.8 | sht, spk, nec | 2.9 | gal, sht, spk, nec | 3.8 | gal, spk |
| C42G | CR291 | 3.5 | stm, spk, spt | 2.1 | spk | 3.5 | spk |
| C42M | CR202 | 3.6 | spk | 2.4 | spk | 4.0 | gal, stm, spk |
| C51C | CR076 | 3.2 | spk | 2.1 | spk | 2.7 | spk |
| C66A | LA2137-2 | 2.7 | spk | 2.1 | spk | 4.3 | gal, spk, nec |
| MC005 | LA1420 | 2.0 | spk | 2.0 | spk | 4.0 | gal, sht, spk spt, nec |
| Controls |  | Pst25 |  | DC3000ΔΔ |  | DC3000ΔΔΔ |  |
| Index | Accession | Average Score | Phenotype | Average Score | Phenotype | Average Score | Phenotype |
| C24A | RG-PtoR | 1.7 | spk, chl | 2.9 | stm, spk | 3.2 | stm, spk |
|  | RG-prf3 | 3.5 | spk, nec, chl | 3.3 | stm, spk | 3.9 | gal, spk, spt |
| C26C | Heinz 1706 | 3.5 | spk, spt, nec, chl | 2.7 | spk | 3.5 | spk, nec |
| C50A | Moneymaker | 2.8 | spk, nec, chl | 2.5 | spk | 3.1 | spk, spt, sht |
| CXR | hpBti9 | 3.3 | spk | 3.3 | spk | 3.7 | spk |
| CXS | hpPto | 3.3 | spk | 3.2 | spk | 3.6 | spk |

### Apparent PTI-like, but no NTI-like responses, were observed in the accession screen

We compared the disease scores between the three *Pst* strains to deduce whether resistance to a particular strain, generally scored as a ‘2’ or less, was due to NTI or PTI (Fig. S2a,b). Many accessions exhibited more severe symptoms in response to DC3000ΔΔΔ than to DC3000ΔΔ (Table 2 and Table S3). Since the only difference between these two strains is the flagellin-encoding *fliC* gene this observation suggested these accessions have an enhanced response to flg22, flgII-28 or possibly some other flagellin-derived peptide. While we did find some accessions that initially appeared to have NTI-based resistance against Pst25 (Fig. S2b), follow up experiments indicated that these accessions expressed Pto, and/or these results could not be repeated. No evidence was observed in any of the 216 accessions for natural variation in an NTI-like response to the 25 TTEs that differ between DC3000ΔΔ and Pst25.

### Use of immune response-specific reporter genes supports primarily PTI-like resistance in some accessions

We used previously developed reporter genes (Pombo *et al*., 2014) to test our hypothesis that some accessions had a PTI-like response to *Pst*. A laccase gene (*LAC*, Solyc04g072280) served as an NTI reporter, and genes encoding a NAC domain protein (*NAC*, Solyc02g069960) and an osmotin-like protein (*OLP*, Solyc11g044390) served as reporters for PTI (Pombo *et al*., 2014). Since in most cases it is unknown which of the 216 accessions might have *Pto*, we induced NTI in a manner that did not elicit a Pto-mediated response. For NTI, we compared the relative expression of *LAC* when plants were inoculated with DC3000ΔΔ versus D36E, a strain lacking all known effectors (Wei *et al*., 2015). For PTI responses, we compared the relative expression of *NAC* and *OLP* when plants were inoculated with DC3000ΔΔ versus DC3000ΔΔΔ. We tested five accessions that we predicted to have PTI responses (EA01356 (C01B), Marmande (C12A), CR202 (C42M), LA2137 (C66A), and LA1420 (MC005)) and found that all five of these accessions had an increased relative expression of one or both PTI reporter genes upon PTI induction (Fig. 2). Two of the accessions (C01B and MC005) also had a small but significant induction of the NTI reporter, but for both accessions a PTI reporter was induced more highly than the NTI reporter, and overall accessions tested with reporter genes primarily had an induction of PTI. Together, these data indicate that reporter genes are a useful tool to confirm or elucidate the hypothesized immune response pathway responsible for resistance.

**Fig. 2.**
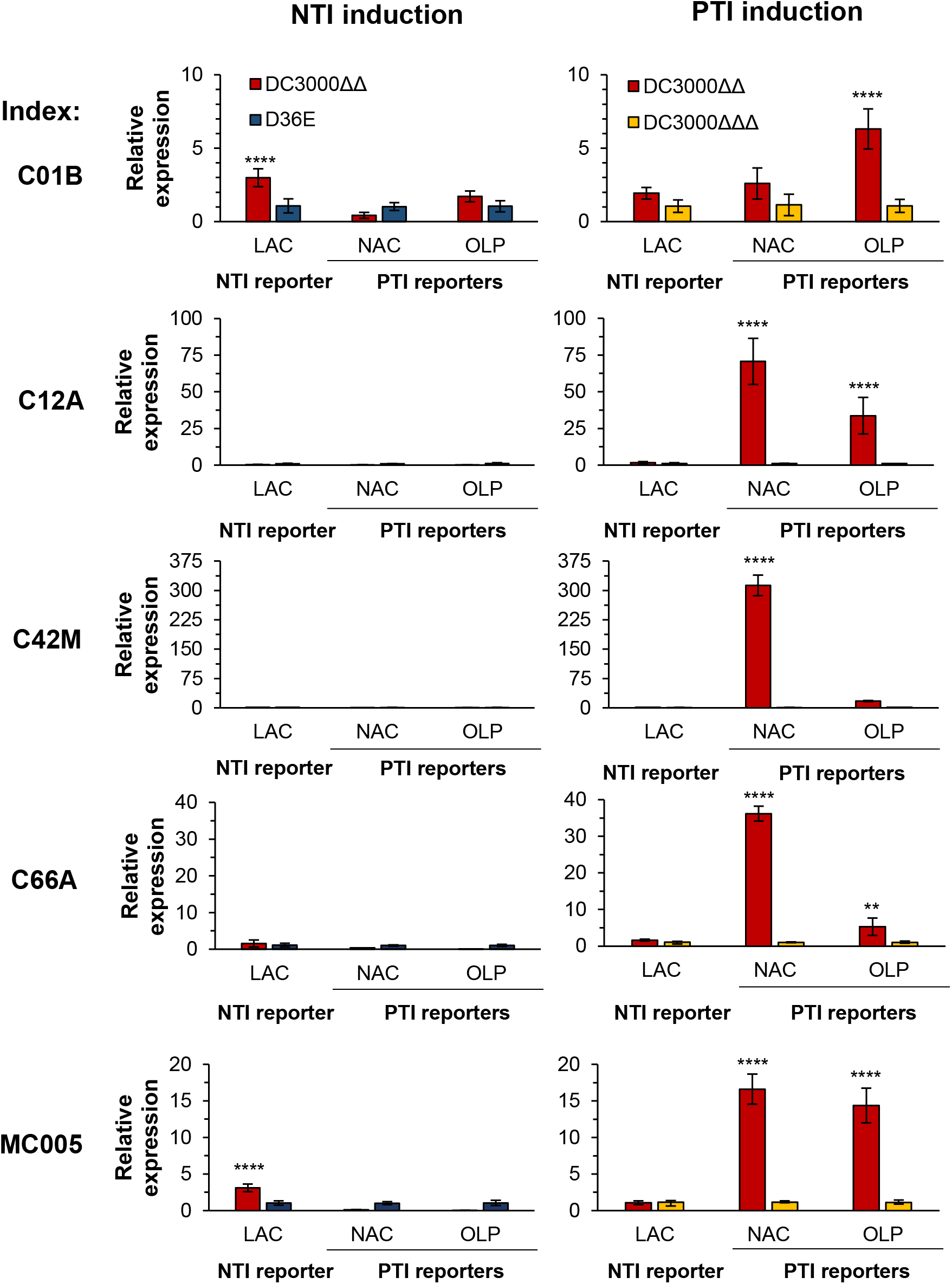
Use of reporter genes to differentiate NTI and PTI responses. Accessions that had hypothesized PTI responses (C01B, C12A, C42M, C66A, and MC005) were tested with reporter genes that are specifically induced during NTI or PTI conditions. NTI induction: DC3000ΔΔ (red bars) vs. D36E (blue bars). PTI induction: DC3000ΔΔ (red bars) vs. DC3000ΔΔΔ (yellow bars). Laccase (LAC, Solyc04g072280) served as an NTI reporter, and a NAC domain protein (NAC, Solyc02g069960) and an osmotin-like protein (OLP, Solyc11g044390) served as reporters for PTI (Pombo *et al*., 2014). Bars indicate the mean of three plants and error bars represent +/− S.D. Significance was determined by a pairwise t-test and significance is indicated with asterisks \*\**P*<0.01, \*\*\*\**P* < 0.0001.

### Leaf surface-based factors play a major role in PTI-mediated resistance to *Pst*

To further characterize the differential PTI-like symptoms observed with the DC3000 strains, we measured bacterial populations in selected accessions. Two accessions were spray inoculated with DC3000ΔΔ and DC3000ΔΔΔ and bacterial populations measured two days later (Fig. 3a). The DC3000 strain lacking flagellin reached a population 150-fold higher than DC3000ΔΔ in these experiments confirming the importance of the PTI response in the tomato-*Pst* interaction. Differential motility of the DC3000 strains with and without flagellin could affect their ability to enter the apoplast so we repeated the experiment on four accessions using vacuum infiltration (Fig. 3b). We again observed statistically significant differences for bacterial populations although the differences were not as great as with spray inoculation (~12-fold). Together, these observations suggest that differences in PTI mechanisms likely exist in the apoplast and at the leaf surface of tomato, such as those that been reported in Arabidopsis (e.g., stomatal closure to prevent pathogen infection (Melotto *et al*., 2006; Zeng *et al*., 2011)).

**Fig. 3.**
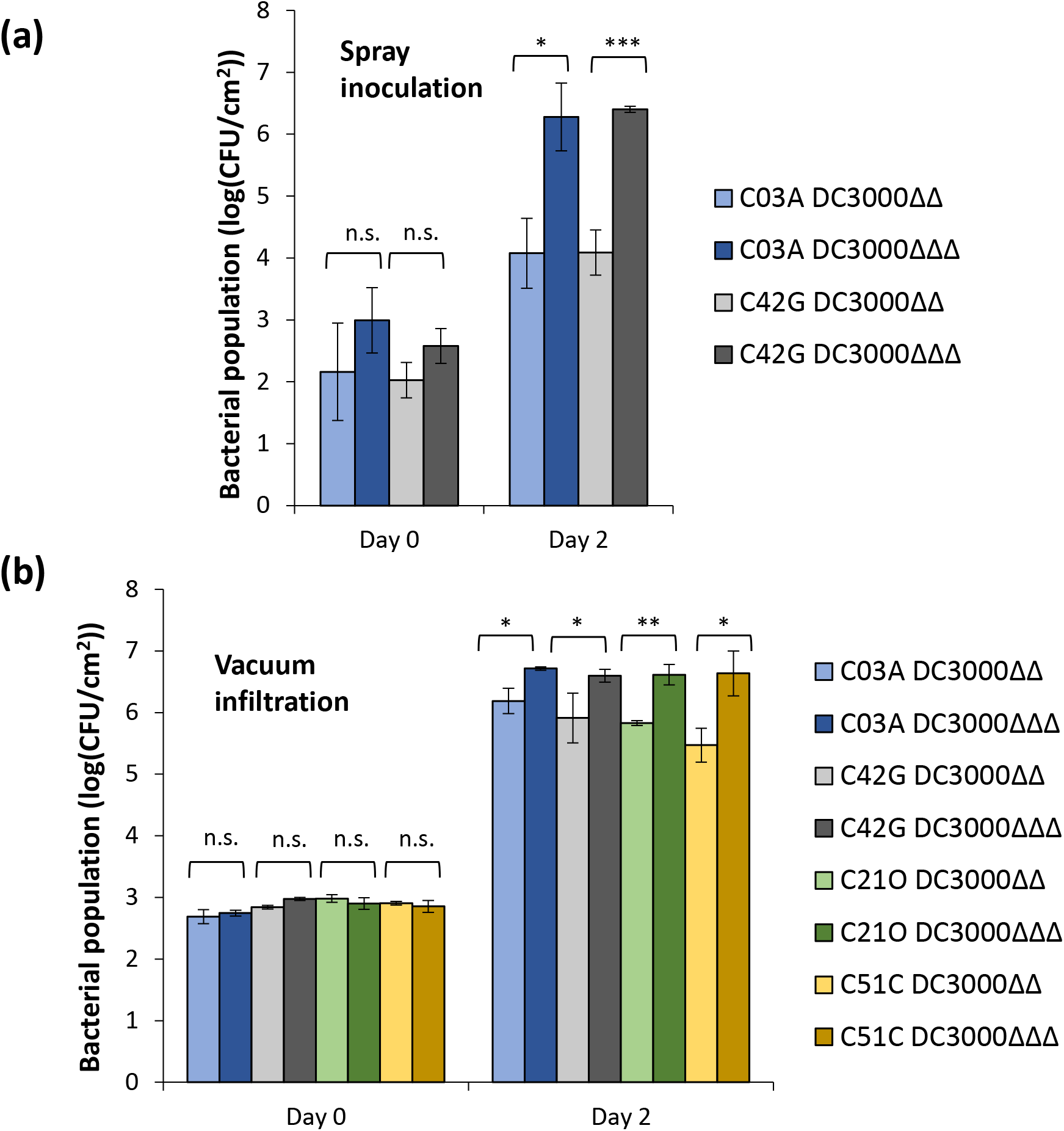
Representative accessions that showed a differential response to strains with and without flagellin. (a) Bacterial populations in leaves after vacuum infiltration of four representative tomato accessions. (b) Bacterial populations in leaves after spray inoculating of two representative accessions. Strains used were DC3000Δ*avrPto*Δ*avrPtoB* (DC3000ΔΔ) and DC3000Δ*avrPto*Δ*avrPtoB*Δ*fliC* (DC3000ΔΔΔ). Significance was determined by a pairwise t-test and significance is indicated with asterisks: \**P*<0.05, \*\**P*<0.01, \*\*\**P*<0.001, not significant (n.s.) P>0.05. Both (a) and (b) are representative of three independent experiments. Bars indicate the mean of three plants and error bars represent +/− S.D.

### Several accessions exhibit divergent responses to flagellin-derived peptides in ROS assays

To test whether the accessions identified as having a PTI-like response in the screen have differential responses to flagellin peptides, we performed reactive oxygen species (ROS) bioassays on fifty-eight accessions using flg22 and flgII-28 peptides (Fig. 4 and Table S7). Rio Grande-PtoR and Heinz 1706 were used as ‘average response’ controls, Yellow Pear as a negative control for flgII-28 (Hind *et al*., 2016) and a CRISPR line knocked out for FLS2 as a negative control for flg22 (Jacobs *et al*., 2017). These experiments revealed accessions that had enhanced responses to flg22 or flgII-28 and in some cases no response to flgII-28 (Fig. 4 and Table S7). All of the fifty-eight accessions responded to flg22, although some responded weakly. Many of the weak responders to flg22 were of *S. pimpinellifolium* background and they also responded weakly to flgII-28, and it is therefore possible that the assay is not optimized for this species. In total, we identified five accessions that had an enhanced response to flg22, four accessions that had an enhanced response to flgII-28, and three accessions that had no response to flgII-28. Accessions that were high responders to one or both peptides were also scored as having less severe disease symptoms after spray inoculation with DC3000ΔΔ. For example, Panama (C13D, average score 2.6), EA01356 (C13E, average score 2.5), OH7983 (C32D, average score 3.3), CR149 (C41C, average score 2.9), CR056 (C42I, average score 2.15), and PI 65023 (C49A, average score 2.4) were all high responders to one or both peptides (Tables S3 and S7). Three accessions, including EA01356 (C13E), had strong responses to both peptides, while CR149 (C41C) had a strong flg22 response but no flgII-28 response. EA01356 (C01B) had an average response to flg22 and no response to flgII-28. Many accessions, for example Marmande (C12A), were identified that have average responses to both peptides. Together, these data suggest that many of the accessions that show some resistance to *Pst* have stronger flagellin-mediated PTI responses, with some of the accessions having novel responses to specific flagellin peptides.

**Fig. 4.**
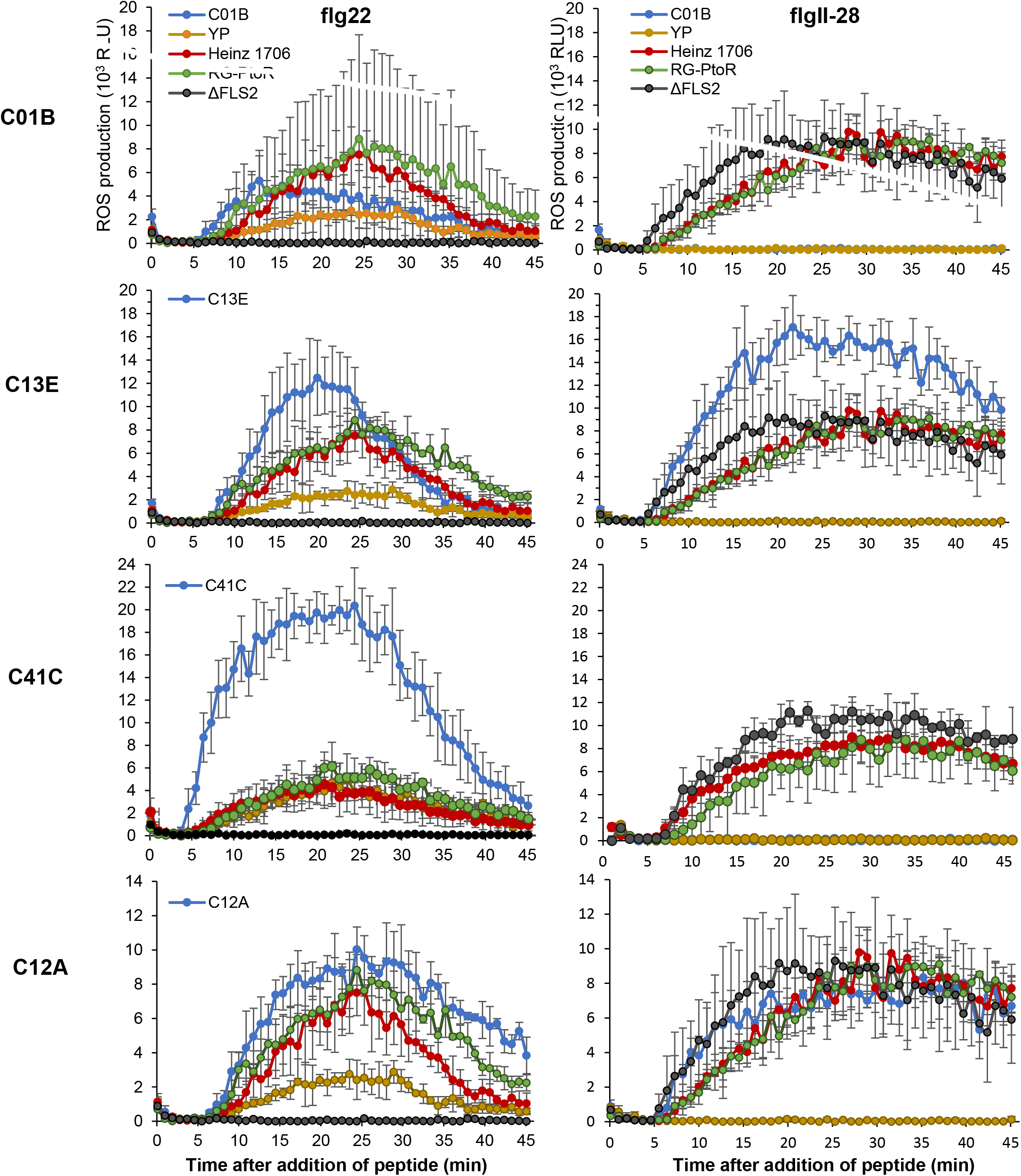
Production of reactive oxygen species in response to flagellin-derived peptides in representative accessions. Reactive oxygen species (ROS) assays were performed on 58 accessions using 100 nM flg22 or 100 nM flgII-28 peptides, and four representative accessions are shown. Bars indicate the mean of at least two plants and error bars represent mean +/− S.D. C01B: average flg22 response, no flgII-28 response. C13E: higher than average flg22 and flgII-28 responses. C41C: higher than average flg22 response, no flgII-28 response. C12A: average responses to both peptides. Heinz 1706 and Rio Grande-PtoR served as ‘average’ response controls. Yellow Pear (YP) served as a negative control for the flgII-28 response. A CRISPR-Cas9 knockout of FLS2 (ΔFLS2) in tomato variety M82 (Jacobs *et al*., 2017) served as a negative control for the flg22 response.

We next tested whether the ROS responses correlated with *Pst* populations in leaves. Using Rio Grande-PtoR as a control, we spray inoculated the four representative accessions previously tested for natural variation in ROS responses to flg22 and flgII-28 (in Fig. 4) and measured bacterial growth of DC3000ΔΔ versus DC3000ΔΔΔ (Fig. 5a). Although we observed increased bacterial growth when *fliC* was deleted, indicating the recognition of flagellin and induction of PTI, accessions that had enhanced responses to flg22 and/or flgII-28 did not support significantly smaller populations of DC3000ΔΔ compared to average responders (Fig. 5a). To test whether there was a difference in spatial recognition of *Pst* that may correlate with ROS, we vacuum infiltrated the same four accessions and measured bacterial growth (Fig. 5b). Again, we found no correlation with the ROS responses and bacterial populations. However, as in Fig. 4 there was again a difference in bacterial growth between spray and vacuum inoculation. While there was a significant difference in bacterial populations between the DC3000 strains with and without flagellin for all the accessions tested by spray inoculation (Fig. 5a), there was a significant difference in populations for only three of the accessions (C01B, C13E, and RG-PtoR) inoculated by vacuum infiltration (Fig. 5b). This suggests that for the other two accessions, C12A and C41C, while flagellin is strongly recognized on the leaf surface, flagellin might be recognized to a lesser extent in the apoplast.

**Fig. 5.**
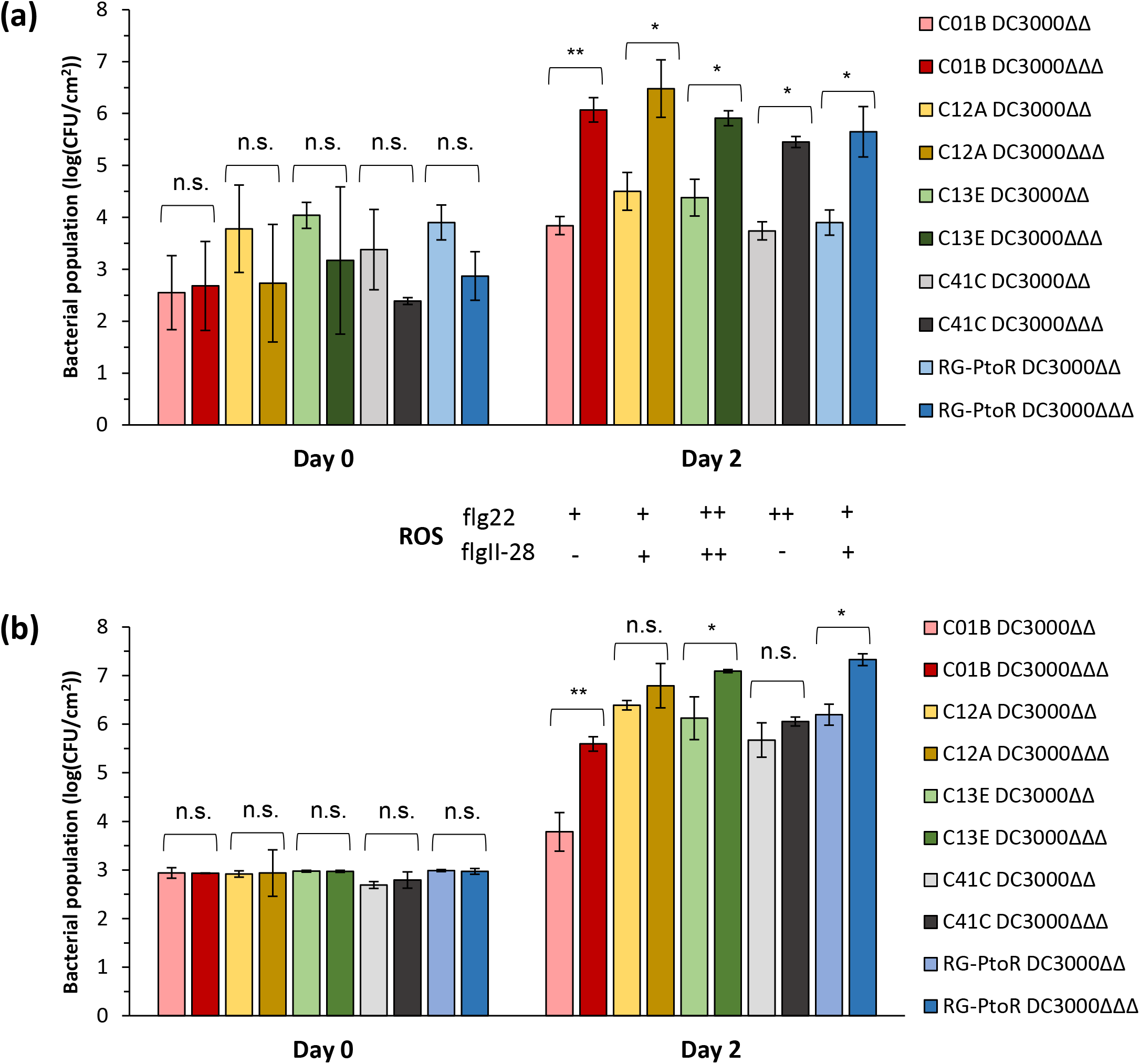
Bacterial population sizes do not correlate with ROS responses. Bacterial populations in leaves after (a) spray inoculation or (b) vacuum inoculation of accessions from Figure 4. Relative ROS responses from accessions in Fig. 4 are displayed below for comparison (−, no response; +, ‘average’ response; ++, ‘high’ response). Strains used were DC3000Δ*avrPto*Δ*avrPtoB* (DC3000ΔΔ) and DC3000Δ*avrPto*Δ*avrPtoB*Δ*fliC* (DC3000ΔΔΔ). Rio Grande-PtoR (RG-PtoR) was used as a control. Significance was determined by a pairwise t-test and significance is indicated with asterisks: \**P*<0.05, \*\**P*<0.01, not significant (n.s.) *P*>0.05. Bars indicate the mean of three plants and error bars represent +/− S.D. The experiment was repeated two (a) or three (b) times with similar results and the figure is representative of the independent experiments.

### Stem gall is a simply inherited trait

To begin the characterization of one of the more unusual phenotypes we examined the genetic basis of the formation of stem galls on some accessions. In the screen such galls were observed primarily on accessions belonging to *S. pimpinellifolium* (mainly clusters 66-83) and upon inoculation with DC3000 strains but not Pst25. In a subsequent inoculation of an additional twenty *S. pimpinellifolium* accessions with DC3000ΔΔ, we observed that nine accessions formed galls and eleven did not (Table S8). Yellow Pear does not form galls while LA1589, a *S. pimpinellifolium* accession, does form galls. We therefore inoculated 156 F2 plants derived from a cross between Yellow Pear × LA1589 (Hind *et al*., 2016) with DC3000ΔΔ and scored them for gall formation. A total of 36 plants formed galls and 120 did not. A chi-square test, where the null hypothesis was that galls did not segregate in a 3:1 ratio, failed to reject the null hypothesis (X^2^ = 0.158, *P*-value > 0.05). Thus, the stem gall phenotype appears to be a recessive trait due to a single locus.

To determine whether viable *Pst* cells are present inside the galls we spray inoculated plants of two gall-forming accessions (C74A and CXG) with DC3000ΔΔ, surface-sterilized the galls with 10% bleach, macerated the tissue, and plated the liquid on King’s B medium amended with rifampicin to select for DC3000ΔΔ. Bacteria grew from both samples within two days. Imaging of a cross section of the galls revealed a clear demarcation between the green stem and the brown gall tissue containing the *Pst* cells (Fig. 6a).

**Fig. 6.**
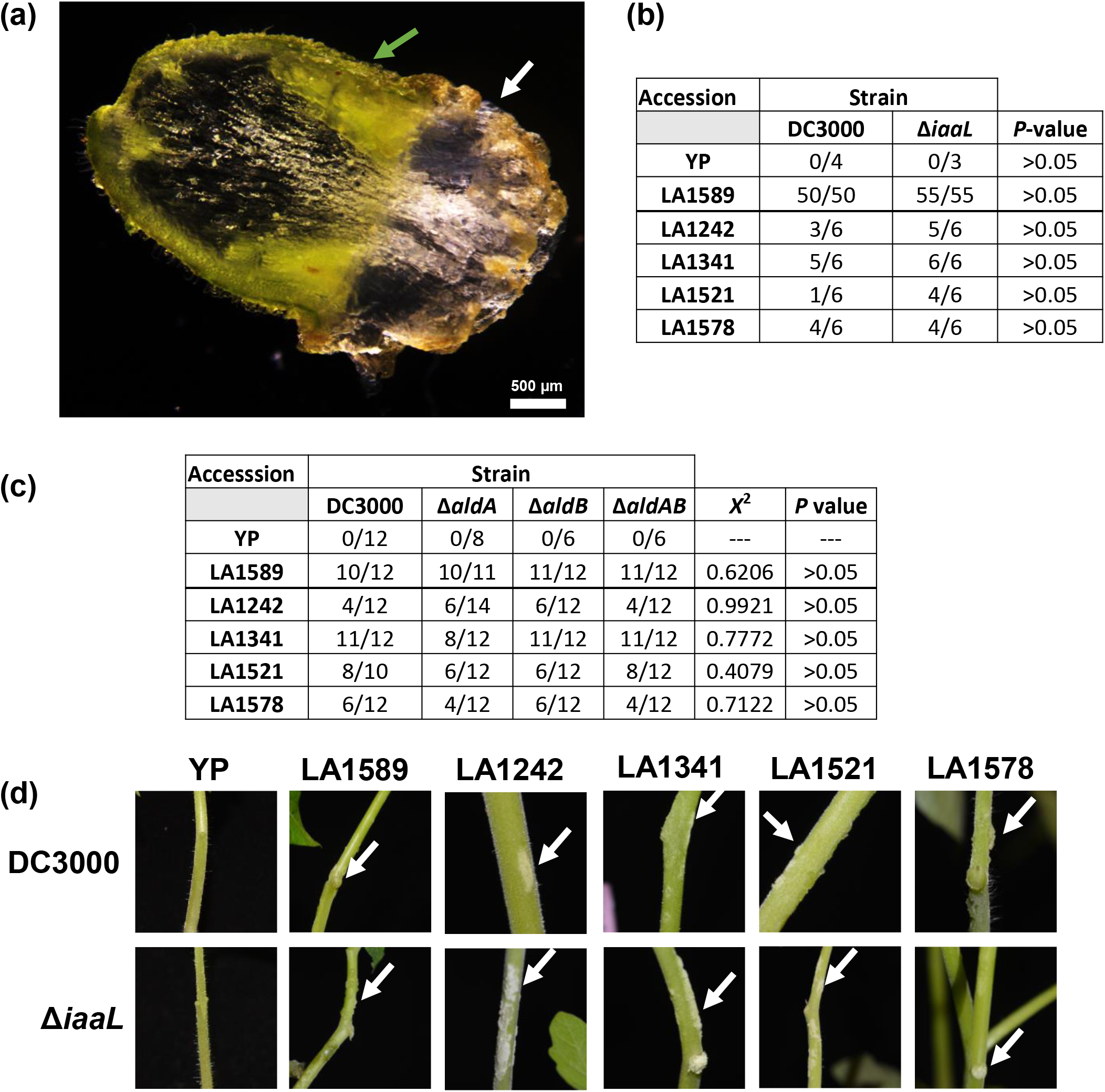
*Pst* strains carrying mutations in three genes affecting *Pst* auxin production still cause gall formation. (a) Cross section image of a gall (brown area marked with a white arrow) that developed on a stem (marked with a green arrow) on accession LA1589 after spray inoculation with DC3000. Scale bar represents 500 μm. (b) Five *S. pimpinellifolium* accessions that form galls were spray inoculated with DC3000 or DC3000Δ*iaaL* (Δ*iaaL*) and gall formation was observed. Shown is the number of plants observed with galls over the total number of plants from three experiments. A Fisher’s exact test was performed to test whether there was a statistically significant difference in gall formation when plants were inoculated with DC3000 versus the mutant strain. (c) Five *S. pimpinellifolium* accessions that form galls were spray-inoculated with DC3000, DC3000Δ*aldA* (Δ*aldA*), DC3000Δ*aldB* (Δ*aldB*), or DC3000Δ*aldA*Δ*aldB* (Δ*aldAB*), and gall formation was observed. Shown is the number of plants observed with galls over the total number of plants from two replicates. A chi-square test was conducted to test whether there was a statistically significant difference in gall formation when plants were inoculated with DC3000 versus the mutant strains. (d) Images of plants sprayed with DC3000 or Δ*iaaL*. White arrows indicate galls. There was no difference observed in gall formation between these two strains. In all experiments, Yellow Pear (YP) served as a negative control that does not form galls. Galls were scored 3 days post inoculation. In (b) and (c), the bold line separates the F2 population parents (YP and LA1589) from the bold line separates the F2 population parents (YP and LA1589) from the other tested *S. pimpinellifolium* accessions.

*Pst* DC3000 produces auxin which plays a role in bacterial pathogenesis (Rosebrock & Caudy, 2017; McClerklin *et al*., 2018) and we hypothesized that this plant hormone might be involved in the formation of the stem galls as it is in other plant-pathogen interactions (Glass & Kosuge, 1986; Glass & Kosuge, 1988; Rosebrock & Caudy, 2017; McClerklin *et al*., 2018). We spray inoculated five gall-forming accessions (LA1589, LA1242, LA1341, LA1521, and LA1578) with four different DC3000 strains that reduce indole-3-acidic acid (IAA) production: DC3000Δ*aldA*, DC3000Δ*aldB*, DC3000Δ*aldA*Δ*aldB*, and DC3000Δ*iaaL* (Lam *et al*., 2014; McClerklin *et al*., 2018). No difference in gall formation was observed in plants inoculated with any of these strains compared to wildtype DC3000 (Fig. 6b,c,d). Thus, we found no evidence that auxin produced by DC3000 is involved in the stem gall formation observed in these *S. pimpinellifolium* accessions.

## Discussion

Molecular mechanisms that underlie plant resistance to bacterial speck disease have been intensively studied, although such studies typically used just one *Pst* strain and only a few tomato genotypes. By strategically using three *Pst* strains that allow for inferences about the type of immunity and over 200 genetically diverse accessions, we discovered remarkable natural variation for the host response to *Pst* in tomato. Some of these responses involve flagellin-mediated PTI, but many other unusual host phenotypes were observed that could be due to either a host defense or susceptibility. This screen and the subsequent experiments demonstrate that tomato has diverse strategies to respond to *Pst* and therefore lays the foundation for advancing our broader knowledge of the diverse molecular mechanisms underlying immunity in plants.

Many unusual host responses not typically associated with *Pst* were observed in the screen (Fig. 1, Table 2, and Tables S4-S6). This unexpected discovery may have been revealed as a result of the large number and genetic diversity of the screened accessions, the more natural (spray) inoculation method, and/or the high humidity conditions used to aid pathogen infection and disease development. While we do not yet know whether these responses are related to resistance or susceptibility, these symptoms could give us insight into the molecular mechanisms of symptom development in the future. For example, screens of *Arabidopsis* mutants identified enhanced disease susceptibility (*EDS*) genes that have been implicated in a number of disease resistance pathways, including the TIR-NLR Roq1 (Recognition of XopQ1) which recognizes the *P. syringae* effector HopQ1 and activates NTI (Reuber *et al*., 1998; Volko *et al*., 1998; Schultink *et al*., 2017). It is possible that some of the unusual symptoms we observed represent natural enhanced disease susceptibility mutations that could be leveraged to discover novel molecular mechanisms governing resistance. It is also possible that some of the unusual symptoms are resistance responses that could directly be mapped and integrated into future breeding programs.

One unusual phenotype, ‘stem gall’, was observed primarily with *S. pimpinellifolium* lines, and in an F_2_ population derived from a tomato and *S. pimpinellifolium* cross we observed a 3:1 segregation for the stem gall phenotype that suggests a recessive, single-locus trait. We also identified a few tomato accessions that formed galls in the screen, and it will be interesting to investigate whether these accessions share a common introgressed segment from *S. pimpinellifolium*. We have yet to determine whether gall formation is associated with a distinct ‘resistant’ or ‘susceptible’ phenotype because galls develop on accessions with both limited and advanced foliar symptoms (Table S3-S6). However, we have isolated bacteria from surface-sterilized galls which suggests that bacteria enter and proliferate in the gall tissue. Interestingly, we only observed gall formation when plants were spray inoculated in high humidity (as opposed to vacuum infiltrated). Because auxin is known to play a role in formation of galls on woody hosts (Glass & Kosuge, 1988) and is important for the virulence of DC3000 (Mutka *et al*., 2013; Kunkel & Harper, 2018; McClerklin *et al*., 2018), we investigated whether auxin plays a role in *Pst* stem gall formation. DC3000 strains with mutations in auxin biosynthetic genes still induced gall formation; however, these strains still produce some auxin as the *ΔiaaL* mutant only reduces lysine-conjugated IAA not free IAA (Glass & Kosuge, 1988), and the *ΔaldAB* double mutant still accumulates some auxin (McClerklin *et al*., 2018). Therefore, at this stage we cannot entirely rule out a role of bacteria-derived auxin in stem gall formation. When we compared these auxin-related genes present in DC3000 and Pst25, we found that these genes are all highly conserved between the two *Pst* strains. Compared to DC3000, the AldA protein sequence varies by one amino acid in Pst25 (E258Q), and AldB varies by two amino acids (G238S, V445L), while the iaaL protein is 100% identical between the two strains. Since DC3000 and not Pst25 induces gall formation, this also supports the idea that bacteria-derived auxin may not be responsible for gall production. Further comparisons of the Pst25 and DC3000 genome sequences are needed to identify other potential gall-related genes.

*Pto* is the only known source of NTI-mediated *Pst* resistance in tomato and it originates from the closely-related wild relative *S. pimpinellifolium*. Our screen was designed to represent diversity across cultivated tomato and *S. pimpinellifolium* to search for new sources of resistance to *Pst*. While we found many accessions with unique PTI responses, we did not identify any accessions with a clear NTI-like phenotype. Our only evidence for potential NTI comes from the use of reporter genes in Fig. 2, which showed a small but significant increase in NTI-related reporter genes in two accessions (C01B and MC005). While it is unknown if this is biologically significant, subsequent vacuum infiltration and bacterial growth assays of one of the accessions (C01B) showed that DC3000ΔΔΔ grew about 25-fold more than DC3000ΔΔ, suggesting that PTI is the major contributor of resistance in C01B (Fig. 5). For the reporter gene assay, we also observed severe disease symptoms with MC005 3 dpi after vacuum infiltration with DC3000ΔΔΔ but saw limited disease for DC3000ΔΔ, suggesting that PTI is also the major contributor to resistance in MC005. This work, and previous screens aimed at finding new sources of NTI-based resistance (Pitblado & Kerr, 1980; Pilowsky, 1982), indicate that no NTI-based resistance to *Pst* is present in cultivated tomato. Comparatively, in a recent screen of 1,041 Arabidopsis accessions using *Pst* DC3000, six accessions displayed an HR-like response to DC3000. Two of these recognized AvrPto and two recognized HopAM1; two others had a weak cell death response (Velasquez *et al*., 2017). The authors suggested that since *Arabidopsis* is unlikely to have co-evolved with *Pst* DC3000, some of the effectors it recognizes might be present in other adapted pathogens and thus *Arabidopsis* has evolved the ability to recognize them and in turn have resistance against DC3000. In the future, it could be useful to screen accessions of the twelve wild species of tomato which might be better candidates for having NTI genes since they are more likely to have co-evolved with *Pst*.

We identified some accessions that produced more ROS than average accessions in response to flg22 and/or flII-28 and also showed reduced symptoms, especially under natural infection conditions where the DC3000 strain lacking flagellin grew about 150-fold more than the strain expressing flagellin (Fig 3b). Two accessions, C03A and C42G, had few symptoms when inoculated with DC3000ΔΔ but were more susceptible to DC3000ΔΔΔ (Table S3). While leaf morphology may play a role in resistance (e.g., waxy cuticles, dense trichomes, and complex leaf topography), we showed that the recognition of flagellin is the major resistance determinant in the accessions tested in Fig 3b. Interestingly, while previous studies have found PTI results in a moderate inhibition of pathogen growth (~10-fold) (Zipfel *et al*., 2004; Lacombe *et al*., 2010; Schwizer *et al*., 2017), our screen revealed a much stronger inhibition of pathogen growth (~150-fold) when plants recognize the pathogen on the leaf surface (Fig. 3), which is more similar to the degree of growth inhibition expected from NTI. This may also be due to a spatial requirement and mechanistic difference when PTI is activated at the leaf surface versus in the apoplast. Two accessions, C12A and C41C, had a significant increase in the growth of DC3000ΔΔΔ compared to DC3000ΔΔ when spray inoculated, but no significant increase in growth when vacuum infiltrated (Fig. 5). It is possible that these two accessions primarily recognize flagellin at the leaf surface, as may be expected in a natural infection, but once bacteria enter the apoplast some other unknown MAMP or bacterial factor is recognized that may contribute to resistance. In the recent screen conducted in *Arabidopsis* (Majoros *et al*., 2017), of the 1,041 accessions tested only two accessions were found to have a “surface-based” mechanism of resistance and completely lost resistance when syringe infiltrated. We show that from a screen of 216 accessions, six accessions recognized flagellin more strongly on the leaf surface, but resistance was still maintained when bacteria entered the apoplast, suggesting distinct mechanisms of resistance in tomato and *Arabidopsis*. Further investigation will be needed to identify and characterize the potential host factors involved in this resistance.

We observed natural variation in PTI for the production of ROS in response to flg22 and flgII-28 (Fig. 4). When we measured *Pst* populations in accessions that vary in their ROS responses, we saw no correlation between ROS production levels and degree of bacterial growth (Fig. 5). This might not be unexpected since ROS production is just one readout of the overall PTI response. A previous study comparing disease in the field to responses to flg22, flgII-28, and the cold shock protein csp2 (recognized by CORE) also found no correlation between disease and peptide recognition (Veluchamy *et al*., 2014). Further research is needed to understand the basis for the differences in ROS production, which could be due to differences in the rate of FLS2/FLS3 receptor turnover, expression levels of the proteins, variation in the protein sequence or, for accessions that are high responders to both peptides, natural variation in an unknown common component of the FLS2/FLS3 signaling pathway. Some accessions, such as C01B, did not respond to flgII-28 and had an ‘average’ response to flg22, but yet had a strong PTI phenotype as indicated by the disease scores (Table S3) and bacterial growth (Fig. 5). For this accession, FLS2 may be sufficient to activate the strong PTI response, or it is possible this accession recognizes another flagellin-associated MAMP as reported recently in kiwi fruit (Ciarroni *et al*., 2018).

We identified accessions with unusual host responses and strong PTI-based resistance, and future identification of the gene(s) involved in these phenotypes could reveal new insights into the plant immune system. Genome-wide association studies (GWAS) approaches could be particularly useful, especially for accessions with distinct ‘plus’ or ‘minus’ phenotypes such as the stem galls. We attempted a GWAS with the 20 *S. pimpinellifolium* accessions we tested for galls (Table S8) but were unsuccessful at finding any gene candidates, likely due to small sample size. Phenotyping more tomato or *S. pimpinellifolium* accessions for galls could increase the power of GWAS. In addition to GWAS, we are developing F_2_ populations to enable a bulked segregant analysis and ‘mapping-by-sequencing’ approach to find gene candidates. Eventually, cloning the gene(s) and determining the molecular mechanisms of resistance could significantly advance our understanding of plant immunity.

## Supporting information

Fig S1 high resolution image

## Acknowledgments

We thank Dani Zamir, Dilip Panthee, Sam Hutton, Martha Mutschler, Jim Giovannoni, David Francis, and Mathilde Causse for providing seeds. We thank Barbara Kunkel for the DC3000Δ*aldA*, Δ*aldB*, and Δ*aldAB* strains and Brian Swingle for the DC3000Δ*iaaL* strain. We thank Noam Eckshtain-Levi for supporting experiments, Sam Wolfe, Nick Glynos and Nicole Avellanet for greenhouse assistance, and Brian Bell, Jay Miller, Brittany Fletcher, and Nick Vail for plant care. Funding was provided by National Science Foundation grant IOS-1546625 (GBM, ARC, SRS). K. S. was partially supported by the China Scholarship Council and the National Natural Science Foundation of China (31822046 and 31772355).

## Author Contributions

R.R., S.M., A.F.P, S.R.S., S.R.H., A.C., and G.B.M. designed research; R.R., S.M., A.F.P., A.E.L., K.S., and S.R.S. performed research; R.R., S.M., A.F.P., A.E.L., S.R.S., K.S., and G.B.M. analyzed the data; R.R. and G.B.M wrote the paper.

## Supporting Information

**Fig. S1** Phylogenetic tree of tomato accessions based on whole genome sequences.

**Fig. S2** Host response severity scale and associated scores after spray inoculation.

**Fig. S3** Photographs of unusual symptoms that were due to high humidity (100%).

**Table S1** Accessions included on the phylogenetic tree (see Excel file ‘Table S1’ for complete table).

**Table S2** Accessions used in the screen (see Excel file ‘Table S2’ for complete table).

**Table S3** Scores of all tomato accessions tested in this study for all strains tested (see Excel file ‘Table S3’ for complete table).

**Table S4** Plant phenotypes observed for accessions inoculated with Pst25 (see Excel file ‘Table S4’ for complete table).

**Table S5** Plant phenotypes observed for accessions inoculated with Pst25 DC3000ΔΔ (see Excel file ‘Table S5’ for complete table).

**Table S6** Plant phenotypes observed for accessions inoculated with DC3000ΔΔΔ (see Excel file ‘Table S6’ for complete table).

**Table S7** Reactive oxygen species (ROS) assay data for each accession tested.

**Table S8** Additional *S. pimpinellifolium* accessions tested for the gall phenotype.

## References

Abramovitch RB, Janjusevic R, Stebbins CE, Martin GB. 2006. Type III effector AvrPtoB requires intrinsic E3 ubiquitin ligase activity to suppress plant cell death and immunity. Proc Natl Acad Sci USA 103: 2851–2856.

Abramovitch RB, Kim Y-J, Chen S, Dickman MB, Martin GB. 2003. *Pseudomonas* type III effector AvrPtoB induces plant disease susceptibility by inhibition of host programmed cell death. EMBO J 22: 60–69.

Arredondo CR, Davis RM. 2000. First report of *Pseudomonas syringae* pv. tomato race 1 on tomato in California. Plant Disease 84: 371.

Bao Z, Meng F, Strickler SR, Dunham DM, Munkvold KR, Martin GB. 2015. Identification of a candidate gene in *Solanum habrochaites* for resistance to a race 1 strain of *Pseudomonas syringae* pv. tomato. The Plant Genome 8: 10.3835/plantgenome2015.3802.0006.

Buell CR, Joardar V, Lindeberg M, Selengut J, Paulsen IT, Gwinn ML, Dodson RJ, Deboy RT, Durkin AS, Kolonay JF, et al. 2003. The complete genome sequence of the *Arabidopsis* and tomato pathogen *Pseudomonas syringae* pv. tomato DC3000. Proc Natl Acad Sci USA 100(18): 10181–10186.

Cai RM, Lewis J, Yan SC, Liu HJ, Clarke CR, Campanile F, Almeida NF, Studholme DJ, Lindeberg M, Schneider D, et al. 2011. The plant pathogen *Pseudomonas syringae* pv. tomato is genetically monomorphic and under strong selection to evade tomato immunity. PLoS Pathog 7: e1002130.

Cheng W, Munkvold KR, Gao H, Mathieu J, Schwizer S, Wang S, Yan YB, Wang J, Martin GB, Chai J. 2011. Structural analysis of *Pseudomonas syringae* AvrPtoB bound to host BAK1 reveals two similar kinase-interacting domains in a type III effector. Cell Host & Microbe 10(6): 616–626.

Chinchilla D, Bauer Z, Regenass M, Boller T, Felix G. 2006. The *Arabidopsis* receptor kinase FLS2 binds flg22 and determines the specificity of flagellin perception. Plant Cell 18: 465–476.

Ciarroni S, Clarke CR, Liu H, Eckshtain-Levi N, Mazzaglia A, Balestra GM, Vinatzer BA. 2018. A recombinant flagellin fragment, which includes the epitopes flg22 and flgII-28, provides a useful tool to study flagellin-triggered immunity. J. Gen. Plant Pathol 84: 169–175.

Cunnac S, Chakravarthy S, Kvitko BH, Russell AB, Martin GB, Collmer A. 2011. Genetic disassembly and combinatorial reassembly identify a minimal functional repertoire of type III effectors in *Pseudomonas syringae*. Proc Natl Acad Sci USA 108: 2975–2980.

Glass NL, Kosuge T. 1986. Cloning of the gene for indoleacetic acid-lysine synthetase from *Pseudomonas syringae subsp. savastanoi*. J Bacteriol 166(2): 598–603.

Glass NL, Kosuge T. 1988. Role of indoleacetic acid-lysine synthetase in regulation of indoleacetic acid pool size and virulence of *Pseudomonas syringae subsp. savastanoi*. J Bacteriol 170(5): 2367–2373.

Gomez-Gomez L, Boller T. 2000. FLS2: an LRR receptor-like kinase involved in the perception of the bacterial elicitor flagellin in *Arabidopsis*. Mol Cell 5(6): 1003–1011.

Hind SR, Strickler SR, Boyle PC, Dunham DM, Bao Z, O’Doherty IM, Baccile JA, Hoki JS, Viox EG, Clarke CR, et al. 2016. Tomato receptor FLAGELLIN-SENSING 3 binds flgII-28 and activates the plant immune system. Nat Plants 2: https://doi.org/10.1038/nplants.2016.1128.

Jacobs TB, Zhang N, Patel D, Martin GB. 2017. Generation of a collection of mutant tomato lines using pooled CRISPR libraries. Plant Physiol 174: 2023–2037.

Janjusevic R, Abramovitch RB, Martin GB, Stebbins CE. 2006. A bacterial inhibitor of host programmed cell death defenses is an E3 ubiquitin ligase. Science 311(5758): 222–226.

Jones JB 1991. Bacterial speck. In: Jones JB, Jones JP, Stall RE, Zitter TA eds. Compendium of tomato diseases. St. Paul, MN: APS Press, 26–27.

Kraus CM, Mazo-Molina C, Smart CD, Martin GB. 2017. *Pseudomonas syringae* pv. tomato strains from New York exhibit virulence attributes intermediate between typical race 0 and race 1 strains. Plant Disease 101: 1442–1448.

Kraus CM, Munkvold KR, Martin GB. 2016. Natural variation in tomato reveals differences in the recognition of AvrPto and AvrPtoB effectors from *Pseudomonas syringae*. Mol Plant 9(5): 639–649.

Krzywinski M, Schein J, Birol I, Connors J, Gascoyne R, Horsman D, Jones SJ, Marra MA. 2009. Circos: an information aesthetic for comparative genomics. Genome Res 19(9): 1639–1645.

Kunkeaw S, Tan S, Coaker G. 2010. Molecular and evolutionary analyses of *Pseudomonas syringae* pv. tomato race 1. Mol Plant-Microbe Interact 23: 415–424.

Kunkel BN, Harper CP. 2018. The roles of auxin during interactions between bacterial plant pathogens and their hosts. J Exp Bot 69(2): 245–254.

Kvitko BH, Park DH, Velasquez AC, Wei C-F, Russell AB, Martin GB, Schneider DJ, Collmer A. 2009. Deletions in the repertoire of *Pseudomonas syringae* pv. *tomato* DC3000 type III secretion effector genes reveal functional overlap among effectors. PLoS Pathog 5: e1000388.

Lacombe S, Rougon-Cardoso A, Sherwood E, Peeters N, Dahlbeck D, van Esse HP, Smoker M, Rallapalli G, Thomma BP, Staskawicz B, et al. 2010. Interfamily transfer of a plant pattern-recognition receptor confers broad-spectrum bacterial resistance. Nat Biotechnol 28(4): 365–369.

Lam HN, Chakravarthy S, Wei HL, BuiNguyen H, Stodghill PV, Collmer A, Swingle BM, Cartinhour SW. 2014. Global analysis of the HrpL regulon in the plant pathogen *Pseudomonas syringae* pv. tomato DC3000 reveals new regulon members with diverse functions. PLoS One 9: e106115.

Li B, Meng X, Shan L, He P. 2016. Transcriptional regulation of pattern-triggered immunity in plants. Cell Host Microbe 19: 641–650.

Lin NC, Abramovitch RB, Kim YJ, Martin GB. 2006. Diverse AvrPtoB homologs from several *Pseudomonas syringae* pathovars elicit Pto-dependent resistance and have similar virulence activities. Appl Environ Microbiol 72(1): 702–712.

Lin NC, Martin GB. 2005. An *avrPto/avrPtoB* mutant of *Pseudomonas syringae* pv. *tomato* DC3000 does not elicit Pto-mediated resistance and is less virulent on tomato. Mol Plant-Microbe Interact 18: 43–51.

Lin T, Zhu G, Zhang J, Xu X, Yu Q, Zheng Z, Zhang Z, Lun Y, Li S, Wang X, et al. 2014. Genomic analyses provide insights into the history of tomato breeding. Nat Genet 46(11): 1220–1226.

Liu X, Grabherr HM, Willmann R, Kolb D, Brunner F, Bertsche U, Kuhner D, Franz-Wachtel M, Amin B, Felix G, et al. 2014. Host-induced bacterial cell wall decomposition mediates pattern-triggered immunity in Arabidopsis. eLife 3: e01990.

Majoros A, Platanitis E, Kernbauer-Holzl E, Rosebrock F, Muller M, Decker T. 2017. Canonical and Non-Canonical Aspects of JAK-STAT Signaling: Lessons from Interferons for Cytokine Responses. Front Immunol 8: 29.

Martin GB, Brommonschenkel SH, Chunwongse J, Frary A, Ganal MW, Spivey R, Wu T, Earle ED, Tanksley SD. 1993. Map-based cloning of a protein kinase gene conferring disease resistance in tomato. Science 262: 1432–1436.

Martin GB, Williams JG, Tanksley SD. 1991. Rapid identification of markers near a *Pseudomonas* resistance gene in tomato using random primers and near-isogenic lines. Proc Natl Acad Sci USA 88: 2336–2340.

Mathieu J, Schwizer S, Martin GB. 2014. Pto kinase binds two domains of AvrPtoB and its proximity to the effector E3 ligase determines if it evades degradation and activates plant immunity. PLoS Pathog 10(7): e1004227.

McClerklin SA, Lee SG, Harper CP, Nwumeh R, Jez JM, Kunkel BN. 2018. Indole-3-acetaldehyde dehydrogenase-dependent auxin synthesis contributes to virulence of *Pseudomonas syringae* strain DC3000. PLoS Pathog 14: e1006811.

Melotto M, Underwood W, Koczan J, Nomura K, He SY. 2006. Plant stomata function in innate immunity against bacterial invasion. Cell 126(5): 969–980.

Monaghan J, Zipfel C. 2012. Plant pattern recognition receptor complexes at the plasma membrane. Curr Opin Plant Biol 15(4): 349–357.

Mutka AM, Fawley S, Tsao T, Kunkel BN. 2013. Auxin promotes susceptibility to *Pseudomonas syringae* via a mechanism independent of suppression of salicylic acid-mediated defenses. Plant J 74(5): 746–754.

Oh C-S, Martin GB. 2011. Effector-triggered immunity mediated by the Pto kinase. Trends in Plant Science 16(3): 132–140.

Pascuzzi PE. 2006. Structure-based functional analyses of Pseudomonas type III effector protein AvrPto and evaulation of putative virulence targets in tomato., Ph.D. thesis, Cornell University Ithaca.

Pedley KF, Martin GB. 2003. Molecular basis of Pto-mediated resistance to bacterial speck disease in tomato. Ann Rev Phytopathol 41: 215–243.

Peralta IE, Spooner DM, Knapp S. 2008. Taxonomy of wild tomatoes and their relatives (Solanum sect. Lycopersicoides, sect. Juglandia, sect. Lycopersicon; Solanaceae). USA: Amer Soc Plant Taxonomoists.

Pilowsky M. 1982. Screening wild tomatoes for resistance to bacterial speck pathogen (*Pseudomonas tomato*). Plant Disease: 46–47.

Pilowsky M, Zutra D. 1986. Reaction of different tomato genotypes to the bacterial speck pathogen (*Pseudomonas syringae* pv. tomato). Phytoparasitica 14: 39–42.

Pitblado RE, Kerr EA. 1980. Resistance to bacterial speck (*Pseudomonas tomato*) in tomato. Acta Horticulturae 100: 379–382.

Pombo MA, Zheng Y, Fernandez-Pozo N, Dunham DM, Fei Z, Martin GB. 2014. Transcriptomic analysis reveals tomato genes whose expression is induced specifically during effector-triggered immunity and identifies the Epk1 protein kinase which is required for the host response to three bacterial effector proteins. Genome Biol 15: 492.

Ragonnet-Cronin M, Hodcroft E, Hue S, Fearnhill E, Delpech V, Brown AJ, Lycett S, Database UHDR. 2013. Automated analysis of phylogenetic clusters. BMC Bioinformatics 14: 317.

Rambaut A. 2007. Figtree, a graphical viewer of phylogenetic trees. http://tree.bio.ed.ac.uk/software/figtree/.

Reuber TL, Plotnikova JM, Dewdney J, Rogers EE, Wood W, Ausubel FM. 1998. Correlation of defense gene induction defects with powdery mildew susceptibility in Arabidopsis enhanced disease susceptibility mutants. Plant J 16(4): 473–485.

Riely BK, Martin GB. 2001. Ancient origin of pathogen recognition specificity conferred by the tomato disease resistance gene Pto. Proc Natl Acad Sci USA 98(4): 2059–2064.

Rose LE, Langley CH, Bernal AJ, Michelmore RW. 2005. Natural variation in the *Pto* pathogen resistance gene within species of wild tomato (Lycopersicon). I. Functional analysis of *Pto* alleles. Genetics 171: 345–357.

Rosebrock AP, Caudy AA. 2017. Metabolite Extraction from Saccharomyces cerevisiae for Liquid Chromatography-Mass Spectrometry. Cold Spring Harb Protoc 2017(9): pdb prot089086.

Rosebrock TR, Zeng L, Brady JJ, Abramovitch RB, Xiao F, Martin GB. 2007. A bacterial E3 ubiquitin ligase targets a host protein kinase to disrupt plant immunity. Nature 448: 370–374.

Rosli HG, Zheng Y, Pombo MA, Zhong S, Bombarely A, Fei Z, Collmer A, Martin GB. 2013. Transcriptomics-based screen for genes induced by flagellin and repressed by pathogen effectors identifies a cell wall-associated kinase involved in plant immunity. Genome Biol 14: R139.

Sade-Feldman M, Jiao YJ, Chen JH, Rooney MS, Barzily-Rokni M, Eliane JP, Bjorgaard SL, Hammond MR, Vitzthum H, Blackmon SM, et al. 2017. Resistance to checkpoint blockade therapy through inactivation of antigen presentation. Nat Commun 8(1): 1136.

Sakatos A, Babunovic GH, Chase MR, Dills A, Leszyk J, Rosebrock T, Bryson B, Fortune SM. 2018. Posttranslational modification of a histone-like protein regulates phenotypic resistance to isoniazid in mycobacteria. Sci Adv 4(5): eaao1478.

Schultink A, Qi T, Lee A, Steinbrenner AD, Staskawicz B. 2017. Roq1 mediates recognition of the *Xanthomonas* and *Pseudomonas* effector proteins XopQ and HopQ1. Plant J 92(5): 787–795.

Schwizer S, Kraus CM, Dunham DM, Zheng Y, Fernandez-Pozo N, Pombo MA, Fei Z, Chakravarthy S, Martin GB. 2017. The tomato kinase Pti1 contributes to production of reactive oxygen species in response to two flagellin-derived peptides and promotes resistance to *Pseudomonas syringae* infection. Mol Plant Microbe Interact 30: 725–738.

Sim SC, Van Deynze A, Stoffel K, Douches DS, Zarka D, Ganal MW, Chetelat RT, Hutton SF, Scott JW, Gardner RG, et al. 2012. High-density SNP genotyping of tomato (*Solanum lycopersicum* L.) reveals patterns of genetic variation due to breeding. PloS one 7(9): e45520.

Sun Y, Li L, Macho AP, Han Z, Hu Z, Zipfel C, Zhou JM, Chai J. 2013. Structural basis for flg22-induced activation of the Arabidopsis FLS2-BAK1 immune complex. Science 342(6158): 624–628.

Tomato Genome Consortium. 2012. The tomato genome sequence provides insights into fleshy fruit evolution. Nature 485(7400): 635–641.

Tomato Genome Sequencing Consortium. 2014. Exploring genetic variation in the tomato (Solanum section Lycopersicon) clade by whole-genome sequencing. Plant J 80(1): 136–148.

Velasquez AC, Oney M, Huot B, Xu S, He SY. 2017. Diverse mechanisms of resistance to *Pseudomonas syringae* in a thousand natural accessions of *Arabidopsis thaliana*. New Phytol 214: 1673–1687.

Veluchamy S, Hind SR, Dunham DM, Martin GB, Panthee DR. 2014. Natural variation for responsiveness to flg22, flgII-28, and csp22 and Pseudomonas syringae pv. tomato in heirloom tomatoes. PLoS One 9(9): e106119.

Volko SM, Boller T, Ausubel FM. 1998. Isolation of new Arabidopsis mutants with enhanced disease susceptibility to Pseudomonas syringae by direct screening. Genetics 149(2): 537–548.

Wang L, Albert M, Einig E, Furst U, Krust D, Felix G. 2016. The pattern-recognition receptor CORE of Solanaceae detects bacterial cold-shock protein. Nat Plants 2: 16185.

Wei HL, Chakravarthy S, Mathieu J, Helmann TC, Stodghill P, Swingle B, Martin GB, Collmer A. 2015. *Pseudomonas syringae* pv. tomato DC3000 type III secretion effector polymutants reveal an interplay between HopAD1 and AvrPtoB. Cell Host Microbe 17: 752–762.

Wei HL, Chakravarthy S, Worley JN, Collmer A. 2013. Consequences of flagellin export through the type III secretion system of *Pseudomonas syringae* reveal a major difference in the innate immune systems of mammals and the model plant *Nicotiana benthamiana*. Cell Microbiol 15(4): 601–618.

Wei HL, Zhang W, Collmer A. 2018. Modular study of the type III effector repertoire in *Pseudomonas syringae* pv. tomato DC3000 reveals a matrix of effector interplay in pathogenesis. Cell Rep 23: 1630–1638.

Worley JN, Russell AB, Wexler AG, Bronstein PA, Kvitko BH, Krasnoff SB, Munkvold KR, Swingle B, Gibson DM, Collmer A. 2013. *Pseudomonas syringae pv. tomato* DC3000 CmaL (PSPTO4723), a DUF1330 family member, is needed to produce L-allo-isoleucine, a precursor for the phytotoxin coronatine. Journal of bacteriology 195: 287–296.

Zeng L, Velasquez AC, Munkvold KR, Zhang J, Martin GB. 2012. A tomato LysM receptor-like kinase promotes immunity and its kinase activity is inhibited by AvrPtoB. Plant J 69: 92–103.

Zeng W, Brutus A, Kremer JM, Withers JC, Gao X, Jones AD, He SY. 2011. A genetic screen reveals *Arabidopsis* stomatal and/or apoplastic defenses against *Pseudomonas syringae* pv. tomato DC3000. PLoS Pathog 7(10): e1002291.

Zipfel C, Robatzek S, Navarro L, Oakeley EJ, Jones JD, Felix G, Boller T. 2004. Bacterial disease resistance in Arabidopsis through flagellin perception. Nature 428(6984): 764–767.

