## Supporting information legends for "Natural variation for unusual host responses and flagellin-mediated immunity against *Pseudomonas syringae* in genetically diverse tomato accessions"

### Figures

**Fig. S1** Phylogenetic tree of tomato accessions based on whole genome sequences. The tree includes 1,429 accessions from SolCap (Blanca *et al.*, 2015), tomato '360' (Lin *et al.*, 2014), and tomato '150' (Aflitos *et al.*, 2014). A total of 6,225 SNPs were used, which is a subset of the SolCap SNPs. There are 83 clusters (shown in different colors). The center track (in blue) shows reference allele (Heinz 1706) frequency in the 1,429 accessions along the tomato chromosomes and unanchored regions (in black).

**Fig. S2** Host response severity scale and associated scores after spray inoculation. (a) Details of the severity scoring scale. (b) Distribution of scores recorded in the first screen using NY15125 (Pst25), DC3000ΔavrPtoΔavrPtoB (DC3000ΔΔ), and DC3000ΔavrPtoΔavrPtoBΔfliC (DC3000ΔΔΔ). (c) Photographs of typical bacterial speck symptoms observed in the screen.

**Fig S2** Photographs of unusual symptoms that were due to high humidity (100%). Symptoms that were observed on some accessions due to high humidity conditions alone without *Pst* inoculation.

### Tables

**Table S1** Accessions included in the phylogenetic tree. Accession names, cluster number on the tree, and the source of the genome sequence data are provided. (See Excel file 'Table S1' for complete table).

**Table S2** Accessions used in the screen**.** Available TS codes and Sequence Read Archive run accession (SRR) numbers and source of the seed is shown. The cluster number refers to the location on the phylogenetic tree, and the index is the unique identifier for accessions in the screen. Short read archive (SRA) number is provided. (See Excel file 'Table S2' for complete table).

**Table S3** Scores of all tomato accessions tested in this study for all strains tested. Scores included are the averages of the first and second replicate for every strain tested. A score of '0' indicates no germination (See Excel file 'Table S3' for complete table).

**Table S4** Plant phenotypes observed for accessions inoculated with Pst25. Symptoms were documented in the second round of screening for all accessions above the thick black line, whereas symptom data for accessions below the thick black line were documented in the first round of screening. No data were collected for gray cells. Controls are highlighted in orange (see Excel file 'Table S4' for complete table).

**Table S5** Plant phenotypes observed for accessions inoculated with DC3000ΔΔ. Symptoms were documented in the second round of screening for all accessions above the thick black line, whereas symptom data for accessions below the thick black line were documented in the first round of screening. No data were collected for gray cells. Controls are highlighted in orange (See Excel file 'Table S5' for complete table).

**Table S6** Plant phenotypes observed for accessions inoculated with DC3000ΔΔΔ. Symptoms were documented in the second round of screening for all accessions above the thick black line, whereas symptom data for accessions below the thick black line were documented in the first round of screening. No data were collected for gray cells. Controls are highlighted in orange (see Excel file 'Table S6' for complete table).

**Table S7** Reactive oxygen species (ROS) assay data for each accession tested. 58 accessions that appeared to have a PTI-like response in the initial screen were tested for their responses to flg22 and flgII-28. Scores are from at least two independent replicates using at least two plants, represented by three leaf discs per plant. Rating scale (based on maximum amplitude of response): no response (not significantly higher than negative control) = 0, low response (significantly lower than Rio Grande) = 1, average response (not significantly different than Rio Grande) = 2, high response (significantly higher than Rio Grande) = 3, very high response (at least twice the amplitude of Rio Grande) = 4. In each experiment, Rio Grande served as a positive control for both flg22 and flgII-28 peptides, Yellow Pear served as a negative control for flgII-28 (Hind *et al.*, 2016), and a CRISPR-Cas9-generated line that has a mutation in *FLS2* served as a negative control for flg22 (Jacobs *et al.*, 2017).

**Table S8** Additional *S. pimpinellifolium* accessions tested for the stem gall phenotype.
