## Supplemental figures for "Natural variation for unusual host responses and flagellin-mediated immunity against *Pseudomonas syringae* in genetically diverse tomato accessions"

**Fig. S1** Phylogenetic tree of tomato accessions based on whole genome sequences.

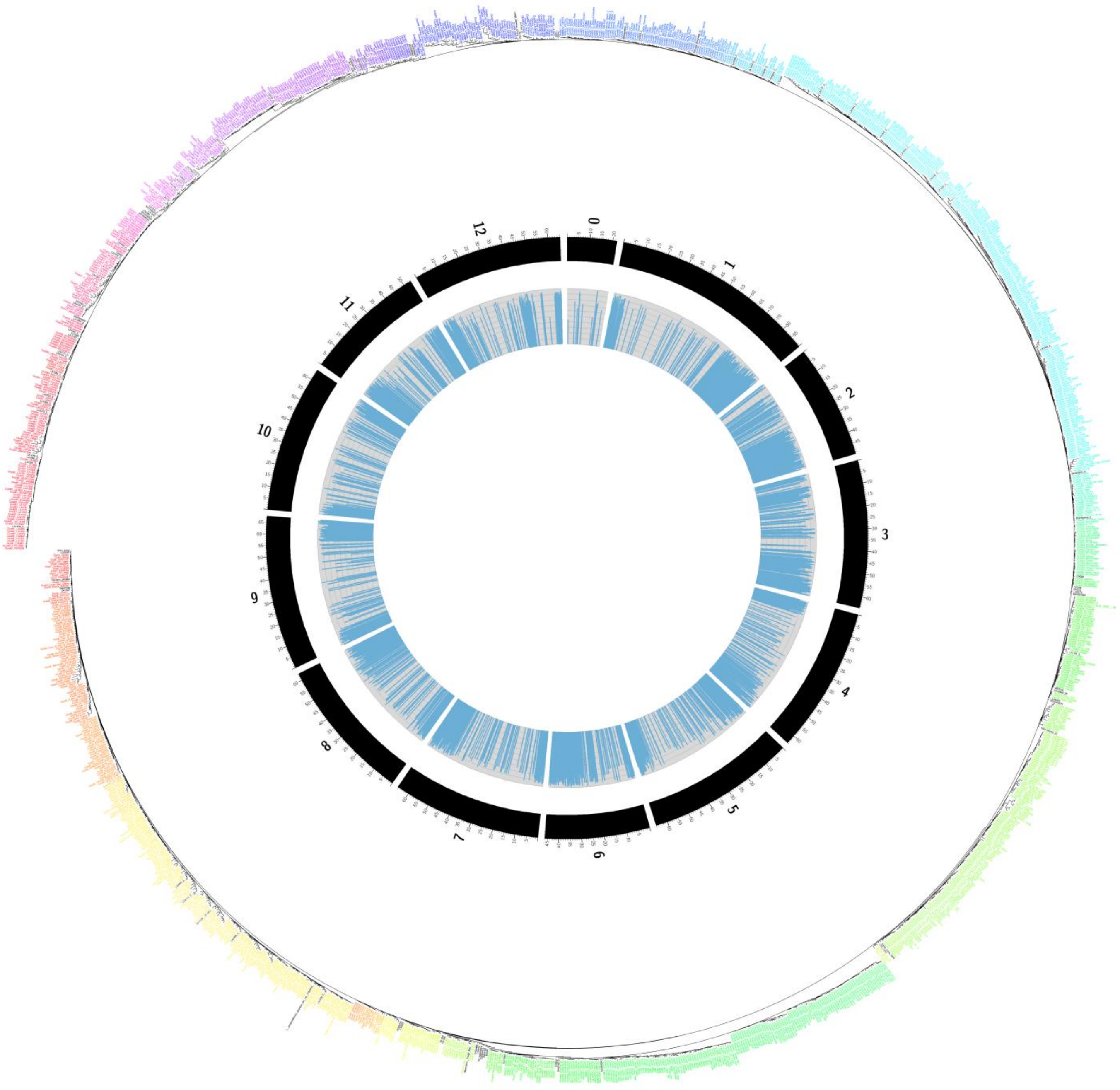


The tree includes 1,429 accessions from SolCap (Blanca *et al.*, 2015), tomato '360' (Lin *et al.*, 2014), and tomato '150' (Aflitos *et al.*, 2014). A total of 6,225 SNPs were used, which is a subset of the SolCap SNPs. There are 83 clusters (shown in different colors). The center track (in blue) shows reference allele (Heinz 1706) frequency in the 1,429 accessions along the tomato chromosomes and unanchored regions (in black).

**Fig. S2** Host response severity scale and associated scores after spray inoculation.

(a)

| Score | Description |
|---|---|
| **1** | No observable symptoms beyond yellow halos (no specks). |
| **2** | A few symptoms on leaves, but limited. No or few symptoms on stem. |
| **3** | Moderate symptoms present on leaves and possibly present on stem. |
| **4** | Severe symptoms on leaves and stem, often accompanied by necrosis and leaf abscission. |
| **5** | Symptoms severe with most or all leaves abscising, leaving only the stem. |

(b)

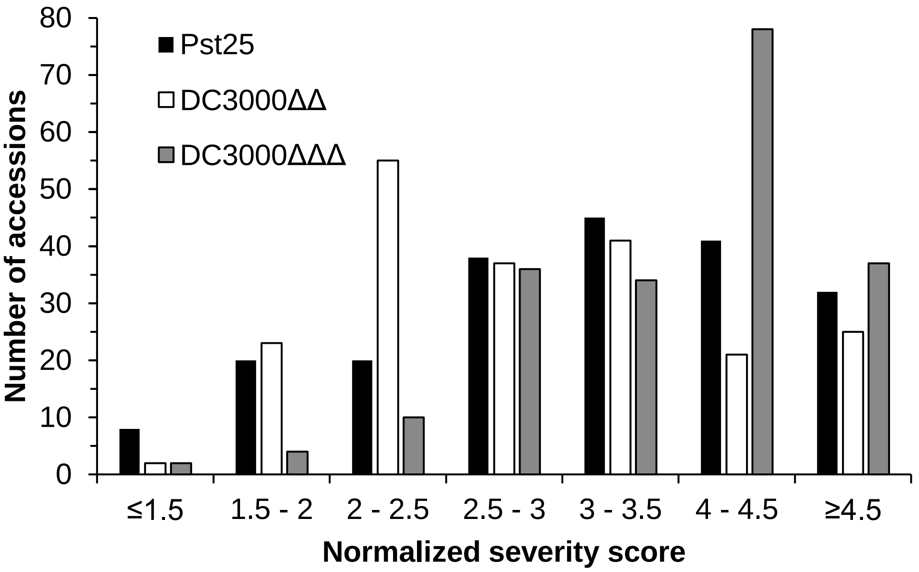


(c)

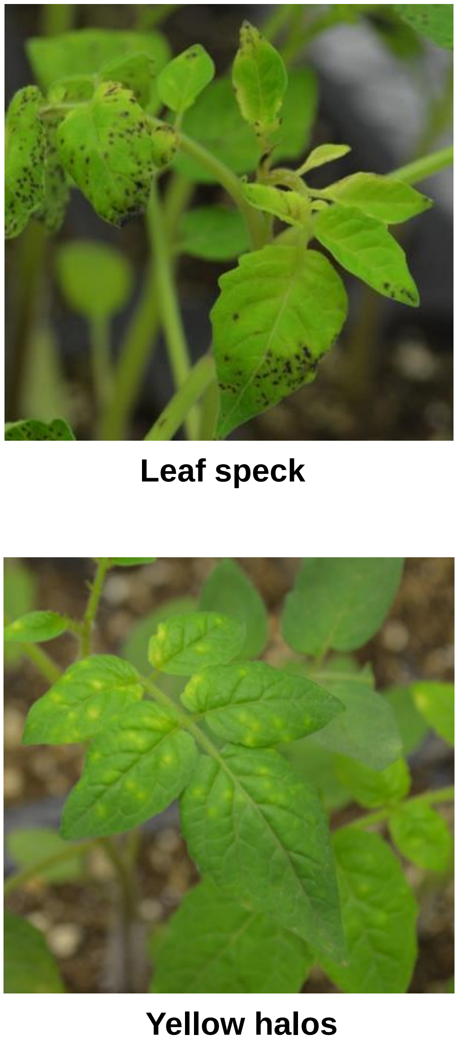


(a) Details of the severity scoring scale. (b) Distribution of scores recorded in the first screen using NY15125 (Pst25), DC3000Δ*avrPto*Δ*avrPtoB* (DC3000ΔΔ), and DC3000Δ*avrPto*Δ*avrPtoB*Δ*fliC* (DC3000ΔΔΔ). (c) Photographs of typical bacterial speck symptoms observed in the screen.

**Fig S3** Photographs of unusual symptoms that were due to high humidity (100%).

**Hypertrophy** **Browning**

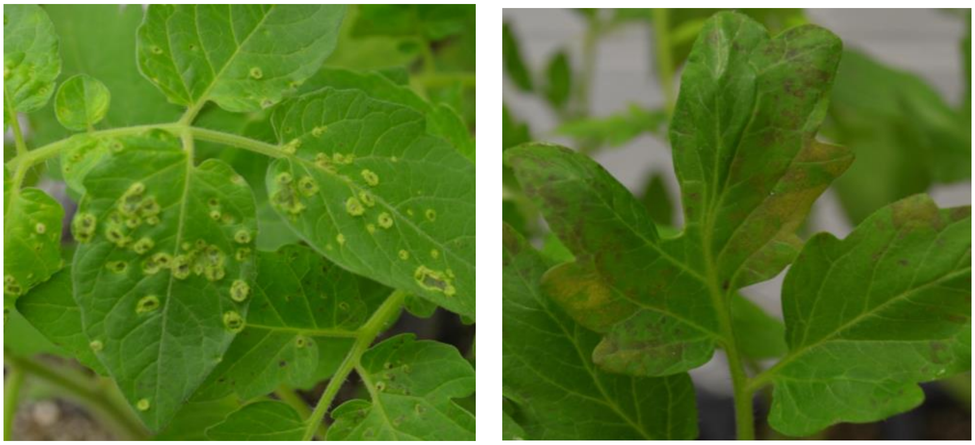

**C67A** **C32C**

Symptoms that were observed on some accessions due to high humidity conditions alone without *Pst* inoculation.

**Figure References**

**Tomato Genome Sequencing Consortium, 2014.** Exploring genetic variation in the tomato (*Solanum* section *Lycopersicon*) clade by whole-genome sequencing. *Plant J* **80**(1): 136-148.

**Blanca J, Montero-Pau J, Sauvage C, Bauchet G, Illa E, Diez MJ, Francis D, Causse M, van der Knaap E, Canizares J. 2015.** Genomic variation in tomato, from wild ancestors to contemporary breeding accessions. *BMC Genomics* **16**: 257.

**Lin T, Zhu G, Zhang J, Xu X, Yu Q, Zheng Z, Zhang Z, Lun Y, Li S, Wang X, et al. 2014.** Genomic analyses provide insights into the history of tomato breeding. *Nat Genet* **46**(11): 1220-1226.
