## Supplementary figures and images for "Natural variation for unusual host responses and flagellin-mediated immunity against *Pseudomonas syringae* in genetically diverse tomato accessions"

### Fig S1 high resolution image

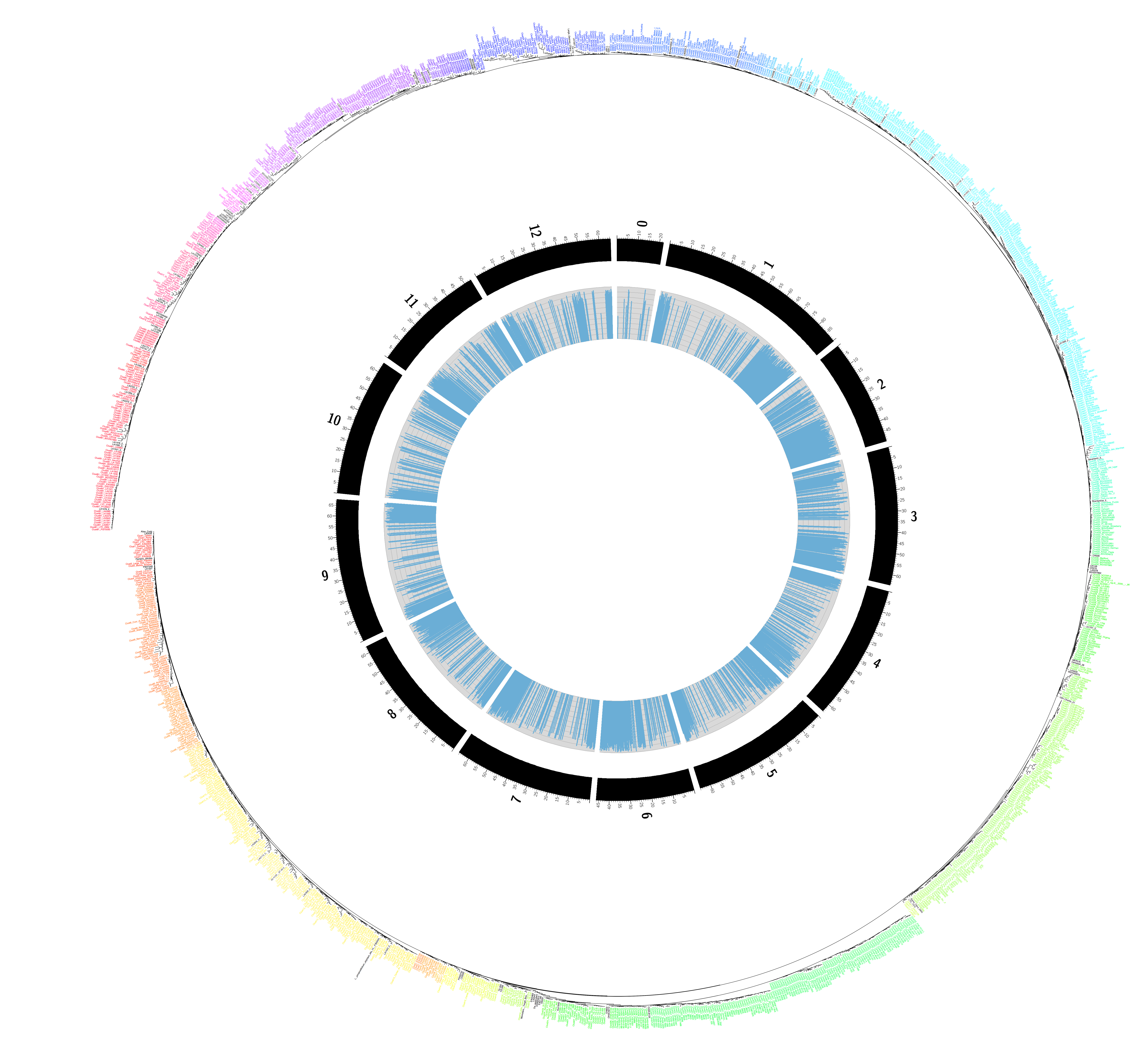
